# Promoter editing of starch branching enzyme IIb and granule-bound starch synthase I balances resistant starch content and amylose content in rice

**DOI:** 10.64898/2026.02.07.704513

**Authors:** YueLong Lin, Qianqi Guo, XingLi Xu, Huaiying Gu, MingHong Hu, Yongjun Wu, Lijun Meng, Guoyou Ye

## Abstract

Increasing attention is being focused on the glycemic index (GI) of daily food for humans, and the resistant starch content (RSC) is an important indicator of GI for starch-rich staple foods. In recent years, some studies revealed that the loss function of single or multiple key enzymes in the primary pathway of starch synthesis substantially increases RSC in rice, such as starch branching enzyme IIb (BEIIb) and soluble starch synthase IIIa (OsSSIIIa). However, a noteworthy negative characteristic of these high RSC mutants is the substantially increased amylose content (AC). AC as a major determinator of rice eating quality, must not be higher than an acceptable limit for most consumers. To solve this problem, in this study, we adopted two promoter editing (PE) editing strategies to develop rice germplasms with a better balance of RSC and AC: one is to edit the promoter of *BEIIb* in a low AC rice variety, another is to edit the promoter of *Waxy* (*Wx*) gene in a *BEIIb* loss of function mutant. Using AC≤20%, which is the range of premium quality rice in China as a criteria, we finally obtained 2 homozygous lines with significantly increased RSC (≥5%) in the NG46 background by promoter editing of *BEIIb* and 1 homozygous line in the YouTang2 (YT2, a *BEIIb* mutant) background by promoter editing of *Wx* gene. Further analysis revealed that AC and the amount of long-chain branches of amylopectin are positively correlated with RSC in the population of *BEIIb* PE lines. However, unexpectedly, the *Wx* PE-line with substantially decreased AC (17.7%) also showed significantly increased RSC (16.9%). Our study not only produces useful germplasms for the high RSC rice breeding in the future but also provides an insight into understanding the relationship between AC and RSC in defective *BEIIb* rice.

## 1. Introduction

Starch is classified into rapidly digestible starch, slowly digestible starch, and resistant starch (RS) based on enzymatic hydrolysis rates in the human digestive system [1]. RS was defined as the fraction of starch that can escape digestion in the upper gastrointestinal tract of healthy individuals, then fermented by the microorganisms in the large intestine to produce some substances beneficial to gut health [2]. According to the differences in source, property, and composition, RS can be further classified into five subtypes, including RS1–RS5 [3, 4]. RS1 is physically protected by cell walls and food matrixes, RS2 is composed of native resistant granules with type B crystallinity, RS3 refers to the retrograded starch formed after cooking, RS4 includes chemically modified or re-polymerized starches by artificial intervention, RS5 belongs to amylose–lipid complexes. Emerging evidence shows that eating starchy foods high in RS content (RSC) is beneficial for blood sugar control, reduces obesity, and improves gut microbiome [5–9]. Therefore, developing high RS foods for humans has been brought into focus by research scholars and breeders recently.

Rice is a staple food for nearly half of the world’s population. Although the starch content in rice grains usually exceeds 70%, their RSC is very low (≤0.5%). Therefore, it’s necessary to develop rice varieties with high RSC. Until now, mutants of some rice genes have been found to have significantly increased RSC, including *BEIIb* [10–12], soluble starch synthase IIa (*SSIIa)* [13], *SSIIIa* [14], and some double-gene combination mutants [15–19]. It seems that there is no shortage of utilizable genes for high RS rice variety development through gene editing. However, these high RS mutants were all accompanied by a dramatic increase in AC and reduced rice eating quality. Thus, searching for more genes or generating quantitative trait variation by editing these known genes are essential to the development of varieties with acceptable RSC and AC at present.

With the rapid advancement of gene editing technology based on the CRISPR-Cas9 system, multiplex gene editing and randomly mutated promoter editing (PE) have become simple and practical. PE was first proposed based on the observation that mutations of cis-regulatory elements (CREs) in gene promoters could generate variants with wide range of phenotypic variation in target trait [20]. It was also found that PE may result in the removal of detrimental pleiotropic effects on nontarget traits of a target gene [20,25,26]. PE has been used for fine manipulation of fruit size in tomato and increasing grain-yield-related traits in maize by multiplex targeting the regulatory region of *SlCLV3* and *CLE* gene, respectively [21, 22]. In rice, PE was applied to create novel *Wx* alleles with fine-tuned amylose levels for better grain quality by editing its promoter or 5’UTR-intron region [23, 24]. Furthermore, introducing mutations in the promoter and 5’-noncoding regions of *OsTB1* created plants with balanced pleiotropic effects of *OsTB1* on different traits, including tiller numbers, plant height, and stem area, enabling them to isolate desirable alleles with more appropriate levels of expression for optimizing breeding targets [25]. Similarly, tiling deletions in *IPA1* cis-regulatory regions also decoupled the negative association between panicle number and size to overcome tradeoffs [26]. These reports showed the magic power of PE not only in creating quantitative trait variations but also in blocking detrimental pleiotropic effects of targeted genes in crops.

Previous studies showed that *BEIIb* mutants in rice significantly increases RSC as well as AC in endosperm in multiple backgrounds. For example, the EM10 mutant in Kinmaze background increase AC from 15.7% to 26.5% [27]. The AC of the ‘Jiangtangdao1’ was increased to about 31.1%, where its wild type ‘Huaqingdao’ was 16.3% [28], and that AC of CRISPR-cas9 edited *BEIIb* mutant in Nipponbare background was about 25% [12]. This level of increase in AC was typically believed to influence taste quality to some extent. In the present study, we aimed at breaking the coupling of high RSC and AC produced by *BEIIb* deficiency to create novel germplasms with high RSC and acceptable AC using two PE strategies by targeting *BEIIb* and *Wx* genes,. First, we design multiplex gRNA cassette targeting the *BEIIb* promoter to create novel alleles with appropriate expression of *BEIIb.* Second, in the same way, multiplex gRNAs were designed to target the promoter of *Wx* in a *BEIIb* mutant rice variety (YT2) to obtain materials with decreased expression of *Wx*. After screening, we finally obtained 4 lines with AC lower than 20% and RSC ranging from 3.8% to 8.4% by *BEIIb* PE and one line with AC (17.7%) and RSC (16.9%) by *Wx* PE, respectively. We found that AC and RSC of *BEIIb* PE lines were positively correlated, however, surprisingly, the *Wx* PE line with substantially lower AC did not decrease RSC. Our study not only produces novel germplasms for high RS rice breedin*g* in future, but also provides some clues for further understanding of the pleiotropic effects of *BEIIb* on RSC and AC.

## 2. Materials and methods

### 2.1 Plant Materials

Rice (*Oryza sativa* L.) ssp. *japonica* Nangeng46 (NG46) and Youtang2 (YT2) were used for the Agrobacterium tumefaciens-mediated transformation. YT2 is an high RSC variety containing the loss function mutant version of *BEIIb* from Jiangtangdao1 [10]. All plant materials were cultivated in the rice transgenic field of Agricultural Genomics Institute at Shenzhen (22°36′22.91″N, 114°29′47.76″E), Shenzhen.

### 2.2 Multiplex gRNAs design and vector construction

To evaluate core regions of a gene promoter for gene expression, data of open chromatin (DNase I hypersensitive sites sequencing, DNase-seq) and conserved elements were collected from PlantRegMap [29], and then 6 sgRNAs were designed by Cas-Designer tools within or surrounding these regions [30]. The vectors of multiplex genome editing were constructed according to the protocol of toolbox [31].

### 2.3 Total RNA extraction and RT–qPCR

Developing seeds of 5 and 10 days after flowering (DAF) were harvested and frozen in liquid nitrogen immediately, and they were then stored at −80℃ until use. Before RNA isolation, developing rice seeds were homogenized using steel balls and a tissue grinder. Total RNA extraction, Reverse transcription, and RT-qPCR were performed as previously described [32].

### 2.4 Measurement of AAC and RS Content

The polished rice was milled to rice flour and passed through a 100-mesh (0.15mm) sieve. AC was measured following the colorimetric method with potassium iodide without the defatting procedure. Briefly, 20±0.1mg rice flour was dispersed in the mixture of 900μL 1M NaOH and 100μL 95% ethanol solution at 85℃ for 20min, during this process, vortexed several times to mix thoroughly. Then these samples were cooled at room temperature(25℃) for 20 minutes, 20μL dispersed starch was added into the mixture of 20μL iodine solution(0.2 g I_2_ and 2 g KI in 100 mL of aqueous solution), 40μL 1M acetic acid, 940μl pure water, then vortexed to mix rapidly and wait for 10 minutes. The absorbance of the solution was measured at 620 nm with a microplate reader, and the standard samples of four levels (0.6%, 8.1%, 14.0%, and 26.9%) were purchased from China National Rice Research Institute. For RS analysis, raw rice flour was measured using a resistant starch assay kit (Megazyme, K-RSTAR) according to the manufacturer’s assay procedure.

### 2.5 Bioinformatics analysis

Coexpression data was obtained by RiceFREND version 2.0 (https://ricefrend.dna.affrc.go.jp/), known transcription factors from the top 2000 of Mutual Rank for *BEIIb* were collected as coexpression gene sets. For prediction of transcription factors binding to *BEIIb* core promoter region, the Gene Search function module of PlantPAN 4.0 (https://plantpan.itps.ncku.edu.tw/plantpan4/) was used. The core region of *BEIIb* promoter from the gRNA target site 2 to 6 with extension of 10 bp on both sides was searched for transcription factor binding sites using the sub-functional module of Promoter Analysis and predictive transcription factors of the list were collected as predictive binding gene sets. The 277 TFs of rice endosperm starch biosynthesis regulatory network were collected according to the study [33].

### 2.6 Dual-Luciferase Reporter Assay

Rice protoplast preparation and transformation were conducted according to the protocol provided by Zhang [34]. About 5μg reporter plasmids and 5μg TF expressed plasmids were co-transformed into 100μL etiolated rice protoplasts. After 16h of dark culture in W5 solution, dual-luciferase activity was measured according to the Dual Luciferase Reporter Gene Assay Kit (Yisheng, Shanghai, China).

### 2.7 Starch Analyses

The microphysical morphology of the endosperm longitudinal section was observed by scanning electron microscopy (HITACHI Regulus 8100, Tokyo, Japan).

For chain length distribution analysis, purified starch (10 mg) was dissolved in 5 mL pure water in a boiling water bath for 60 min. Sodium azide solution (10 μL 2%w/v), acetate buffer (50 μL, 0.6 M, pH 4.4), and isoamylase (10 μL, 1400U) were added to the starch dispersion, and the mixture was incubated in a water bath at 37 ℃ for 24 h. The hydroxyl groups of the debranched glucans were reduced by treatment with 0.5% (w/v) of sodium borohydride under alkaline conditions for 20 h. The preparation of about 600 μL was dried in vacuo at room temperature and allowed to dissolve in 30 μL of 1 M NaOH for 60 min. Then, the solution was diluted with 570 μL of distilled water. The sample extracts were analyzed by high-performance anion-exchange chromatography (HPAEC) on a CarboPac PA-200 anion-exchange column (4.0 * 250 mm; Dionex) using a pulsed amperometric detector (PAD; Dionex ICS 5000 system). Data were acquired on the ICS5000 (Thermo Scientific) and processed using Chromeleon 7.2 CDS (Thermo Scientific).

Gel consistency was evaluated according to GB/T 22294-2008. Triplicate measurements were performed for each sample.

The thermal properties of samples were determined by NETZSCH DSC 300 (Germany) and analyzed as described previously [35].

For X-ray diffraction (XRD) analysis, rice starch samples were passed through a 100-mesh sieve, pressed firmly and spread evenly, then scanned with an X-ray diffraction (Rigaku Miniflex 600 Tokyo, Japan) instrument to obtain the XRD map [36]. The XRD patterns and degree of crystallinity of starch samples were analyzed by Jade 5.0 software.

## 3. Results

### 3.1 Effective promoter editing uses the MGE toolbox to target the promoter core regions

To obtain BEIIb or *Wx* mutants by PE, we designed 6 gRNA expression cassettes (Fig. 1A) targeting the putative core promoter and surrounding region of *BEIIb* and *Wx* promoters, respectively. After screening for homozygous T1 offsprings, the PE lines were sequenced and collected (Fig. 1B). The PE lines of *BEIIb* showed high-frequency of large fragment deletions or insertions (81.8% in all sequenced homozygous mutant plants), with the largest deletion up to 1763bp in PE#22 line. The PE#29 line with complete deletion of the exon1 including coding sequences of *BEIIb* would be considered as a loss of function mutant. In contrast, the *Wx* PE lines showed low frequency (9.1%) of large indels, many mutations were small deletions or insertions near the target sites, only the PE#14 and 24 lines have large fragment deletion between target-1 and target-4, but we detected two large fragments inversion in PE#1-2 and 9-2. These results demonstrated that PE with the MGE toolbox effectively creates mutations on the target sites.

**Fig.1.**
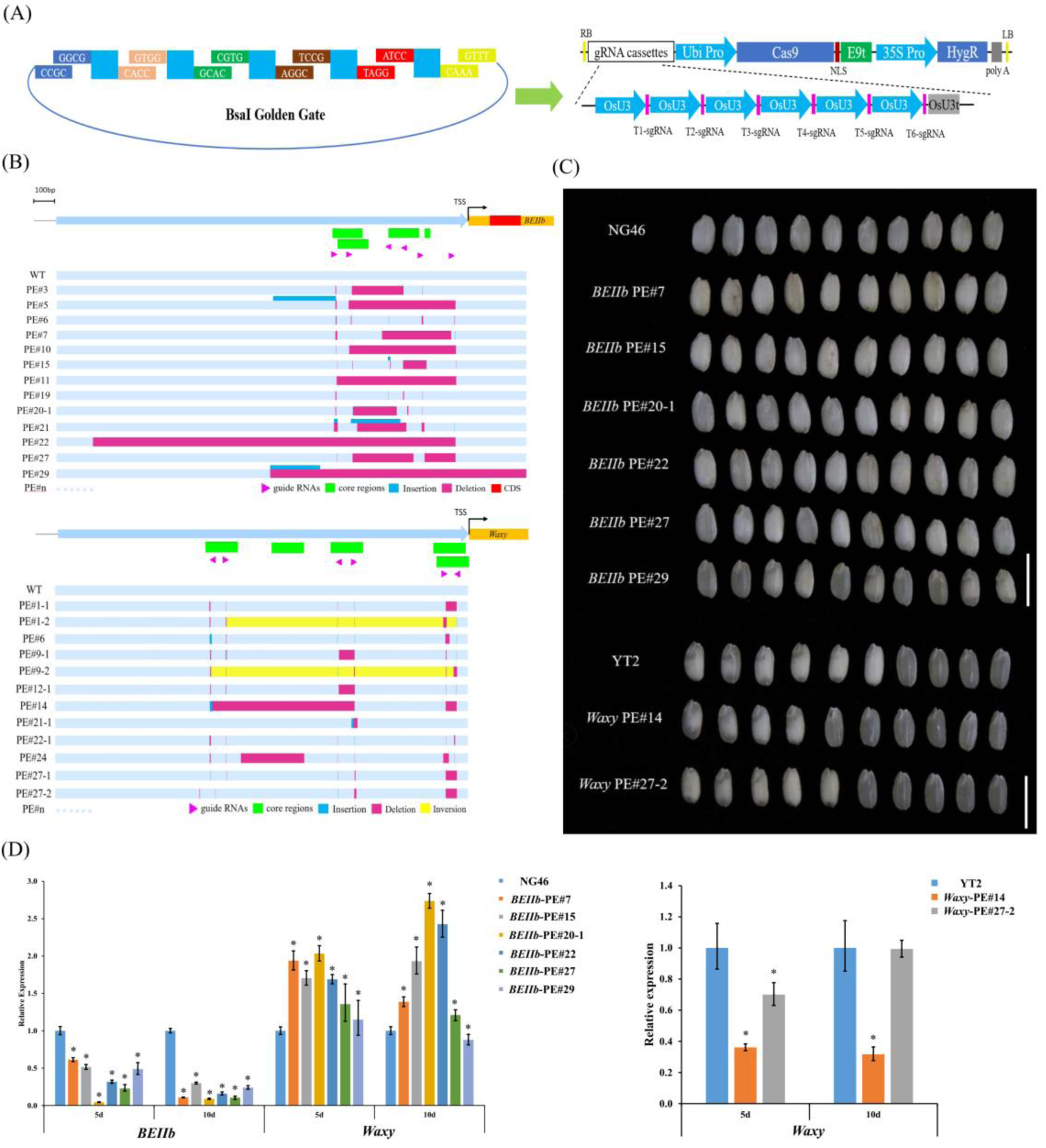
Vector construction, mutation, and expression analysis of promoter editing. (A) The structure diagram of a multiplex-genome-editing vector. (B) Mutation analysis of multiplex gRNA editing for PE lines. (C) Brown rice morphology of PE lines. Bar=1cm. (D) Expression analysis of *BEIIb* or *Wx* gene for PE lines. Bars represent means ± SD, Student’s t-test; * *P* < 0.05

To obtain lines with better balance between RSC and AC. We further conducted selection among the homozygous PE lines based on brown rice appearance and AC. At the end, six *BEIIb* PE lines two *Wx* PE lines were obtained. (Fig. 1C). To confirm the effective mutations of these PE lines to decrease the expression of *BEIIb* or *Wx*, we conducted qRT-PCR using developing seeds of 5 and 10 DAF of these PE lines. Tthe results showed that the expression of *BEIIb* was significantly downregulated in all PE lines, while the expression of *Wx* was significantly upregulated except in the *BEIIb* PE#29 line at 10 DAF (Fig.1 D). We also detected the expression of some other major enzymes in the starch biosynthesis pathway of endosperm, and found that most of these genes showed a significantly increased expression in 5 DAF endosperm including *BEI*, *OsSSI*, *OsSSIIa*, *OsSSIIIa*, *OsSSIIIb* (SP Fig.1). For *Wx* PE lines, *Wx* PE#27-2 line did not show no significantly downregulated expression of *Wx* at 10 DAF (Fig. 1D). Taken together, these results demonstrated the expression of *BEIIb* and *Wx* during rice endosperm development can be drastically decreased by effective promoter editing.

### 3.2 Variations of AC and RSC in PE-lines

To further determine whether the endosperm properties of these PE-lines have changed, we analyzed AC using the Iodine staining method and measured RSC with the Megazyme K-RSTAR Resistant Starch Assay Kit. The results showed that 6 *BEIIb* PE-lines all have significantly higher RSC and AC compared to the WT (Table 1), which is consistent with the expression levels of *Wx* gene in developing endosperm of these six PE lines were also higher than that of WT. In addition, there was a high correlation between AC and RSC within six PE lines (r=0.97) (Fig. 2A). But it is worth noting that when the data of WT with much lower AC and RSC was included, the correlation coefficient was decreased to 0.92 (Fig.2B). Among the six PE lines, *BEIIb* PE#29, which has no entron1 including CDS and is effectively a loss of function mutant, has the highest AC (22.9%) and RSC (17.2%). The relatively lower AC compared to previous studies may be attributed to appropriate promoter editing, but it should be noted that in our study WT was a so-called ‘soft rice’ variety with much lower AC than the varieties used in previous gene editing studies, therefore, this is also an important influencing factor that cannot be ignored completely.

**Fig.2.**
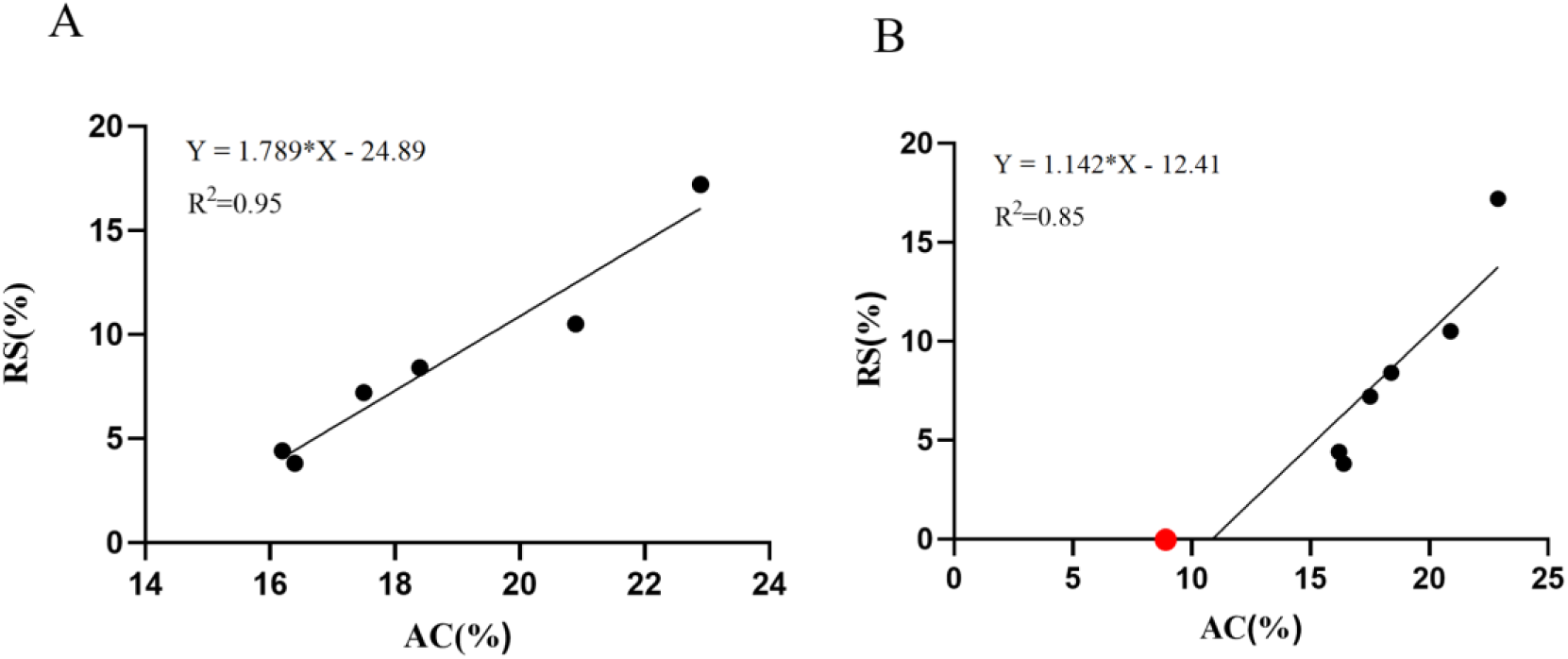
Linear regression analysis between AC and RSC of *BEIIb* PE lines. (A) Linear regression analysis using 6 *BEIIb* PE lines. (B) Linear regression analysis of 6 *BEIIb* PE lines and WT (NG46). The red point represents the WT. Different letters above the bars indicate significant differences (*P* < 0.05) determined by one-way analysis of variance followed by LSD.

To our surprise, the two *Wx* PE lines in the background of a BEIIb loss of function mutant had lower AC but higher RSC compared to WT. PE#14 had the lowest AC (17.7%) but the highest RSC (16.9%) (Table 1). This unexpected result indicated that the commonly observed positive correlation between AC and RSC may be an illusion and could be broken down in selected lines. By downregulating the expression of *Wx* gene through PE, the effect of knocking out BEIIb on AC can be compensated to large extent.

**Table1.**
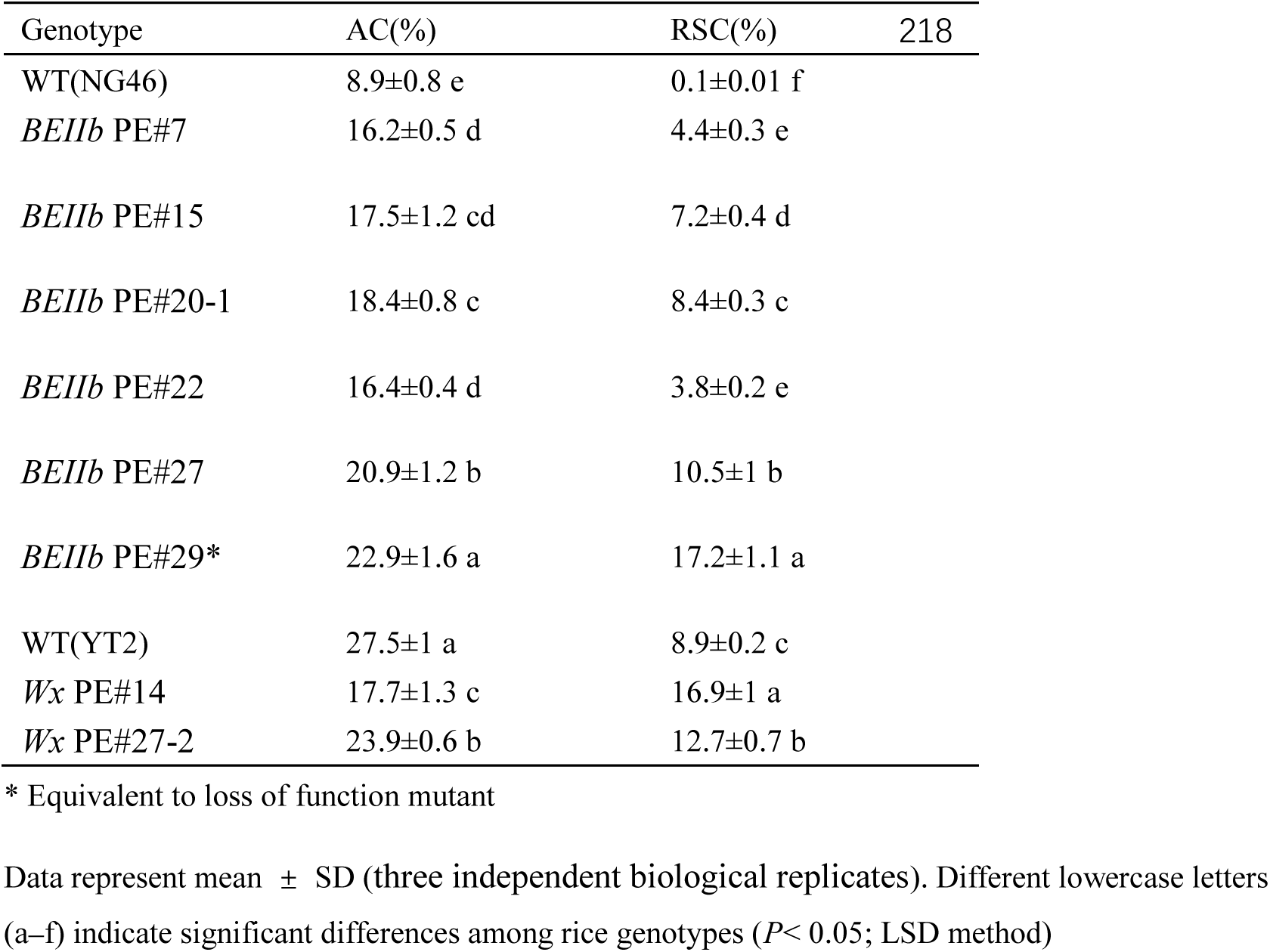

| Genotype | AC(%) | RSC(%) | 218 |
| --- | --- | --- | --- |
| WT(NG46) | 8.9±0.8 e | 0.1±0.01 f |  |
| *BEIIB* PE#7 | 16.2±0.5 d | 4.4±0.3 e |  |
| *BEIIB* PE#15 | 17.5±1.2 cd | 7.2±0.4 d |  |
| *BEIIB* PE#20-1 | 18.4±0.8 c | 8.4±0.3 c |  |
| *BEIIB* PE#22 | 16.4±0.4 d | 3.8±0.2 e |  |
| *BEIIB* PE#27 | 20.9±1.2 b | 10.5±1 b |  |
| *BEIIB* PE#29* | 22.9±1.6 a | 17.2±1.1 a |  |
| WT(YT2) | 27.5±1 a | 8.9±0.2 c |  |
| *Wx* PE#14 | 17.7±1.3 c | 16.9±1 a |  |
| *Wx* PE#27-2 | 23.9±0.6 b | 12.7±0.7 b |  |

\* Equivalent to loss of function mutant
Data represent mean ± SD (three independent biological replicates). Different lowercase letters (a–f) indicate significant differences among rice genotypes ( $P < 0.05$ ; LSD method)

### 3.3 Prediction of key regulatory transcription factors for activation of *BEIIb* in endosperm

Since fragment deletion of the target *BEIIb* promoter region results in its lower expression and thus the high RSC of edited lines, it is attempting to assume that the edited regions (especially from T2 to T6, Fig. 3A) might be the core regulatory region. To this end, we searched for transcription factors (TFs) responsible for activation of *BEIIb* during embryo development. First, we conducted prediction using PlantPAN 4.0 and identified 140 TFs potentially binding to this region. We then used the coexpression database of RiceFREND to analyze TFs coexpression with *BEIIb* and identified 136 TFs. Recently, a rice endosperm starch biosynthesis regulatory network comprised of 277 TFs identified using promoter probes of 15 starch synthesis enzyme-encoding genes (SSEGs) by DNA pull-down combined with LC–MS was constructed [33]. In this network, 17 TFs were for *BEIIb*. However, only *OsERF44* was identified by PlantPAN 4.0 and coexpression analysis and also in the list of 17 TFs confirmed by DNA pull-down. Three TFs including Os07g0182000(*OsbZIP58*), Os03g0410000(*OsMYB3*), and Os02g0710300(*OsRSL2*), were found by both DNA pull-down for binding *BEIIb* promoter and coexpression analysis. In addition, Os08g0490000(*BIM2*), Os07g0644100(*OsbZIP60*), and Os01g0834400(*OsNF-YB2*) identified by both PlantPAN 4.0 and coexpression analysis were among the 277 TFs identified for starch biosynthesis genes.

**Fig.3.**
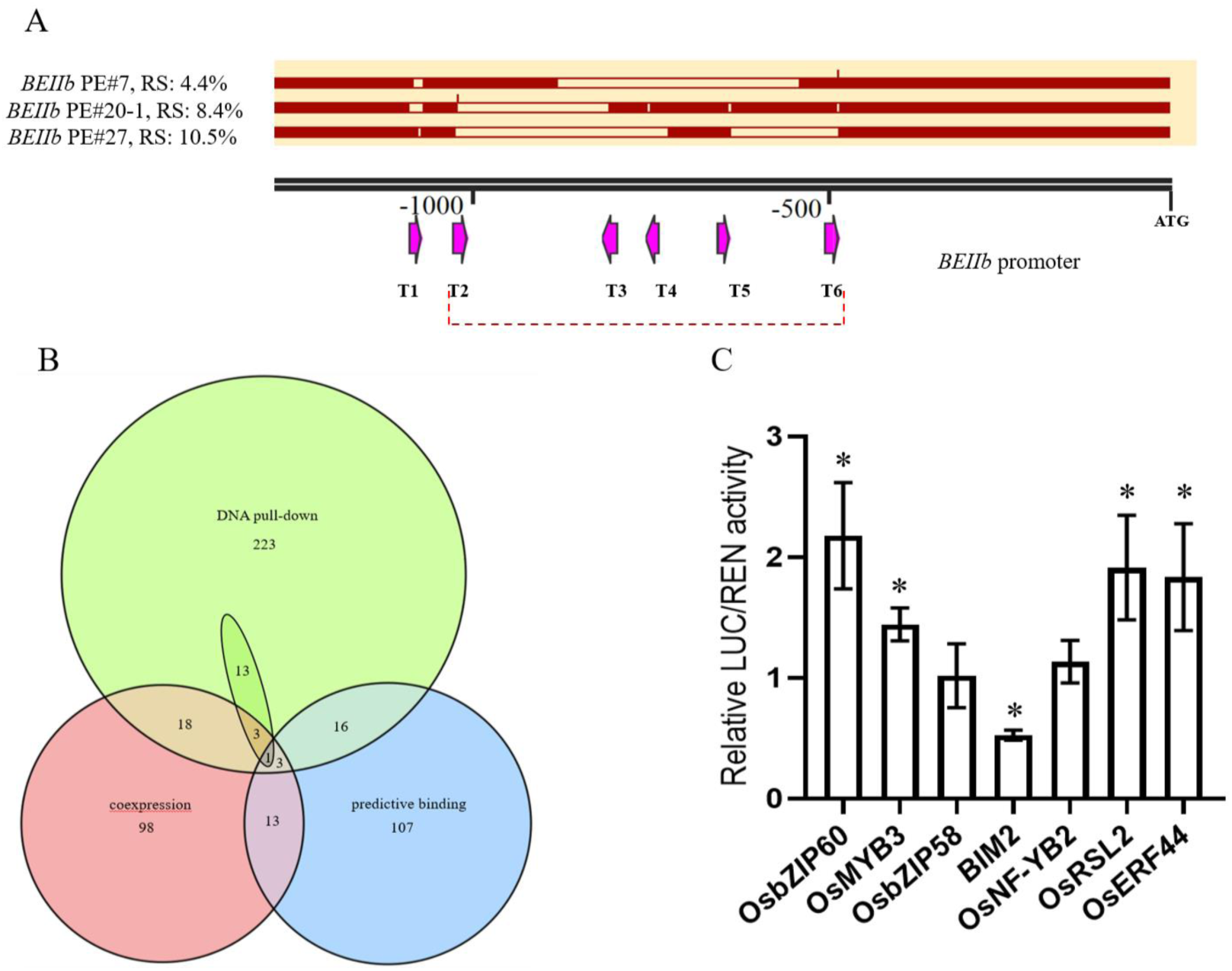
Prediction of transcription factors in the target region of *BEIIb* promoter. (A) Deletion analysis of target region in *BEIIb* promoter. (B) Venn diagram of three gene sets of TFs, the oval box represents the TFs binding the *BEIIb* promoter identified by DNA pull-down. (C) Relative reporter activity (LUC/REN) of target region in rice protoplasts co-expressed with predicted TFs or an empty vector. All data are normalized against the average value expressed empty vector. Bars represent means ± SD (three biological replicates), Student’s t-test; * *P* < 0.05

To further test whether these genes can activate the target region of *BEIIb* promoter, we cloned the sequence of this region into pGreenII0800-LUC vector and then co-transformed with transient expression vectors of these TFs into rice protoplasts to observe the ratio of LUC/REN for empty vector and each of TFs. The results showed that all the four of TFs can activate the expression of *BEIIIb* target region, including *OsbZIP60*, *OsMYB3*, *OsRSL2*, and *OsERF44* (Fig. 3C). Expression of *OsbZIP58* and *OsNF-YB2* in rice protoplast did not significantly activate the expression of *BEIIIb* for the target promoter region, while *BIM2* significantly decreased the expression of *BEIIIb*.

However, due to the limitation of accuracy and coverage for each gene set, and other promoter region may also contribute to the activation of *BEIIb*, since there are still about 5% RS for improvement when compare the *BEIIb* PE#27 and PE#29, we believed that some other TFs may also be important for activation of *BEIIb,* especially those genes within each intersection. Further study should focus on these TFs for regulating the expression of *BEIIb* and try single or multiple gene knockouts to observe the variations of RS production.

### 3.4 The endosperm properties and taste quality of PE-lines are changed

It is well-known that the deficiency of *BEIIb* not only elevated AC and RSC but also changed the morphology of starch granules and the chain length distribution of amylopectin [11, 12, 27, 28]. To test whether our PE-lines showed similar changes and evaluate taste quality, *BEIIb* PE#15, #20-1, and *Wx* PE#14 with acceptable AC (≤ 20%) and RSC ≥ 5% and their WT were mainly selected and tested for starch characteristics and taste quality related parameters. We first scanned the microstructure of starch granules from rice endosperm longitudinal section by electron microscope. As expected, starch granules of the two *BEIIb* PE lines showed spherical and irregularly distributed granules compared to the regular prismatic granules in WT (Fig. 4). Moreover, the lines with higher RSC tends to have increased number of starch granules but with smaller size. Compared to the *be2b* mutant (YT2), *Wx* PE#14 had much fewer irregular fractured gullys on the surface of starch granules (showed by red arrows in Fig. 4) and the starch granules showed multi-spherical protrusions (showed by yellow arrows in Fig. 4). Interestingly, *BEIIb* PE#29 which had the highest RSC among all *BEIIb* PE lines had some starch granules with fractured gullys on the surface, while *BEIIb* PE#15 had big starch granules covered with regular quadrilateral or pentagonal rills (showed by green arrows in Fig. 4).

**Fig.4.**
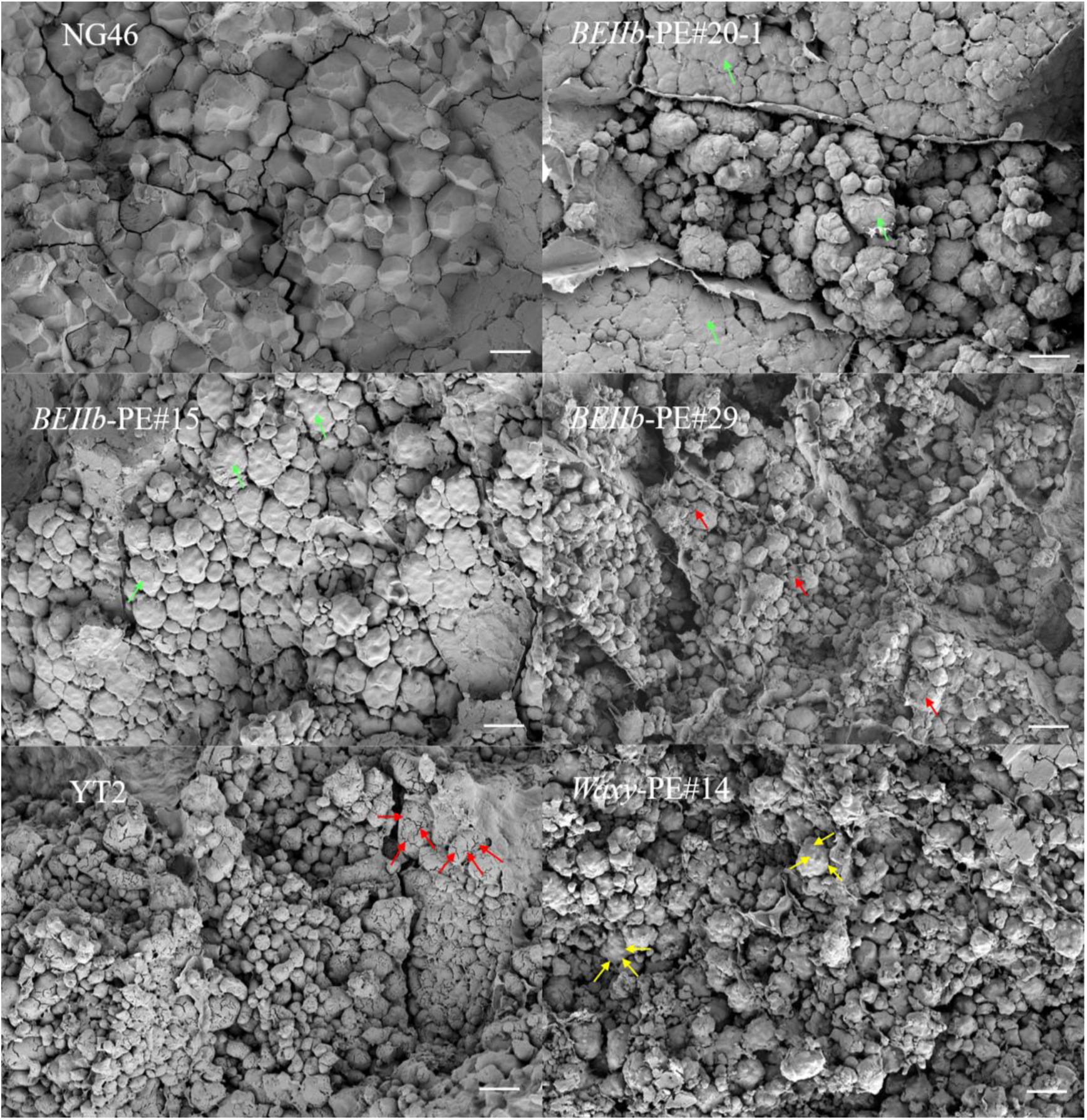
Microphysical structure of endosperm starch granules in PE lines. Bar=10μm

Next, we analyzed chain length distribution of amylopectin for these PE lines. All *BEIIb* PE lines in our analysis showed decrease in short chains DP6-16 and increase in long chains DP18-76 (Fig. 5A). Furthermore, there were three difference peaks at about DP9, DP25, DP50 and two retracement points at about DP17 and DP38, divided chain length distribution of amylopectin into three main zones, from DP6-16, DP18-38 and DP39-76. These results were consistent with previous studies on *be2b* mutant [12, 27]. It is noteworthy that the peak area of short and long chains seems to be associated with the RSC. We test the relationship between total peak area of the changed branching chains and RSC by linear regression analysis and found a highly positive correlation between total long chains (DP18-76) and RSC (r=0.99) and a highly negative correlation between total short chains (DP6-16) and RSC(r=-0.99). This result demonstrated that the RS produced by downregulating *BEIIb* may not be mainly attributed to the enhanced amylose synthesis and the changed branching chains and structural variation of amylopectin is likely more important as previously reported [37].

**Fig.5.**
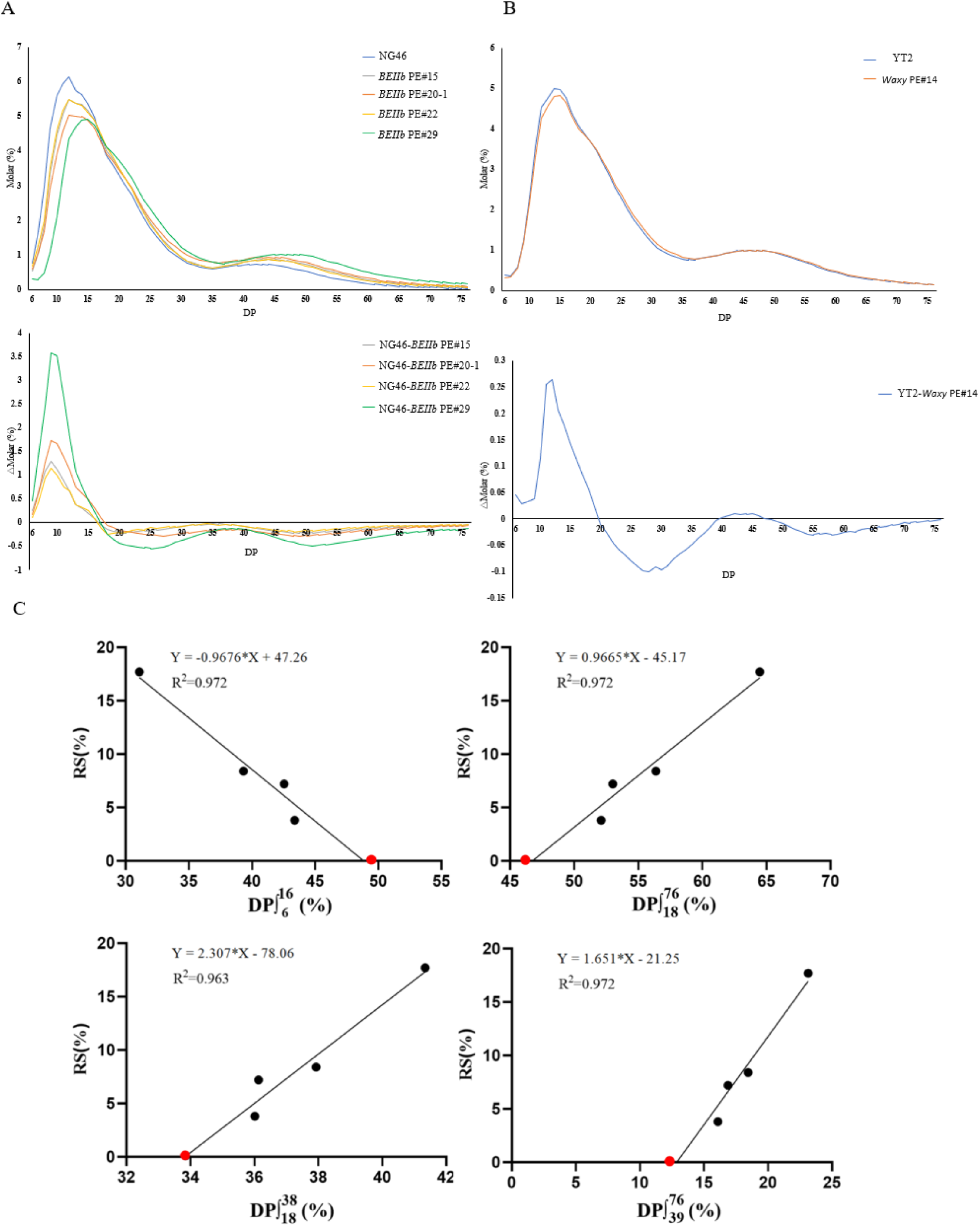
Analysis of amylopectin branching patterns and association with RS content for PE lines. (A) Chain length distribution and differences of endosperm amylopectin for NG46 and *BEIIb* PE lines. (B) Chain length distribution and differences of endosperm amylopectin for YT2 and *Wx* PE#14 line. (C) Linear correlation analysis between different branch chains of amylopectin and RS content in BEIIb PE lines. The red point represents the wild-type NG46 data.

The rice taste quality is affected by many properties of endosperm starch, for example, gelatinization temperature, gel consistency, AC, protein content and lipid content. Among these, AC is an important indicator and is usually negatively correlated with taste quality. In our *BEIIb* PE#15 and PE#20 lines, although their AC increased significantly compared to WT, they were still lower than the 20% standard upper limit of high-quality Japonica rice in China. We also tested gelatinization temperature and gel consistency for these two lines (Table 2), and found that their gel consistency values were > 60mm, which were significantly lower than NG46 (WT) but were still soft (≥60mm), The analysis of thermal properties by Differential Scanning Calorimetry showed that the *BEIIb* PE lines had significantly increased gelatinization temperature. These results demonstrated that lines with high RSC and acceptable eating quality can be obtained by appropriately downregulating *BEIIb* through PE. *Wx* PE#14 line with substantially higher RSC and lower AC than YT2 had significantly higher gel consistency (Fig. 5B), suggesting that it had significantly improved eating taste quality. The higher gelatinization temperature of *Wx* PE#14 line was in line with the more long-chain amylopectin in its endosperm (Fig. 5B). Taken together, these results strongly suggested that PE of BEIIb and *Wx* genes could generate varieties with balanced RSC and taste quality and that simultaneously editing both BEIIb and *Wx* promoters could be a better better good strategy to be further exploited for this purpose.

**Table 2:** Gel consistency and Thermal properties of endosperm starch for PE lines.

| Genotype | Gel consistency<br>(mm) | Thermal properties |  |  |  |
| --- | --- | --- | --- | --- | --- |
|  |  | To (°C) | Tp (°C) | Tc (°C) | ΔH (mJ mg<sup>-1</sup>) |
| WT(NG46) | 111.7±3.1 a | 66.7±3.2 b | 73.3±0.6 c | 77.6±1.9 c | 1.49±0.05 c |
| *BEIIB* PE#15 | 86.7±2.4 b | 72.1±1.2 a | 78.7±1.3 b | 85.1±0.7 ab | 5.9±0.12 a |
| *BEIIB* PE#20-1 | 73±3.6 c | 72.1±3 a | 78.5±0.7 b | 83.9±0.9 b | 2.48±0.02 b |
| *BEIIB* PE#29 | 50.7±2.3 d | 74.2±2.6 a | 81.9±0.6 a | 87.6±2.9 a | 1.03±0.02 d |
| WT(YT2) | 53.7±2.8 b | 68.8±1.6 b | 77.3±2.6 b | 82±1.1 b | 0.29±0.04 b |
| *Wx* PE#14 | 93.7±1.6 a | 78.7±0.8 a | 82±0.9 a | 87.3±0.6 a | 0.67±0.05 a |

### 3.5 Agronomic traits of PE-lines

To evaluate the practicality of these PE-lines for agricultural production, we then investigate some agronomic traits for *BEIIb* PE#15, 20-1, and *Wx* PE#14 in the field. The results were shown in Fig. 6, for plant morphology, plant height, and effective tiller number, all three PE lines all showed no significant difference compared to their wild-type control. Although the *Wx* PE#14 showed a slight but significant decrease in seed setting rate and seed weight, the yield per plant did not significantly decrease in our cultivation conditions, indicating that the total grain numbers of *Wx* PE#14 were more than YT2. For the two *BEIIb* PE lines, the seed weight and yield per plant were significantly decreased with their seed weight being about 96% and 91% of NG46 (WT), respectively. The seed weight of *BEIIb* PE#29, which could be considered as completely loss of function for *BEIIb*, was only about 67% of NG46 (data no showed). The decrease of seed weight and yield of *be2b* mutant was consistent with previous studies. The seed weight of ‘Jiangtagndao1’ was 75% of its original parental variety ‘Huaqingdao’[28], and that of EM10 was 57% of its wild-type plants[16]. The two selected *BEIIb* PE lines, particularly, the *BEIIb* PE#15, had greatly reduced seed weight and yield loss but maintain an acceptable RSC.

**Fig.6.**
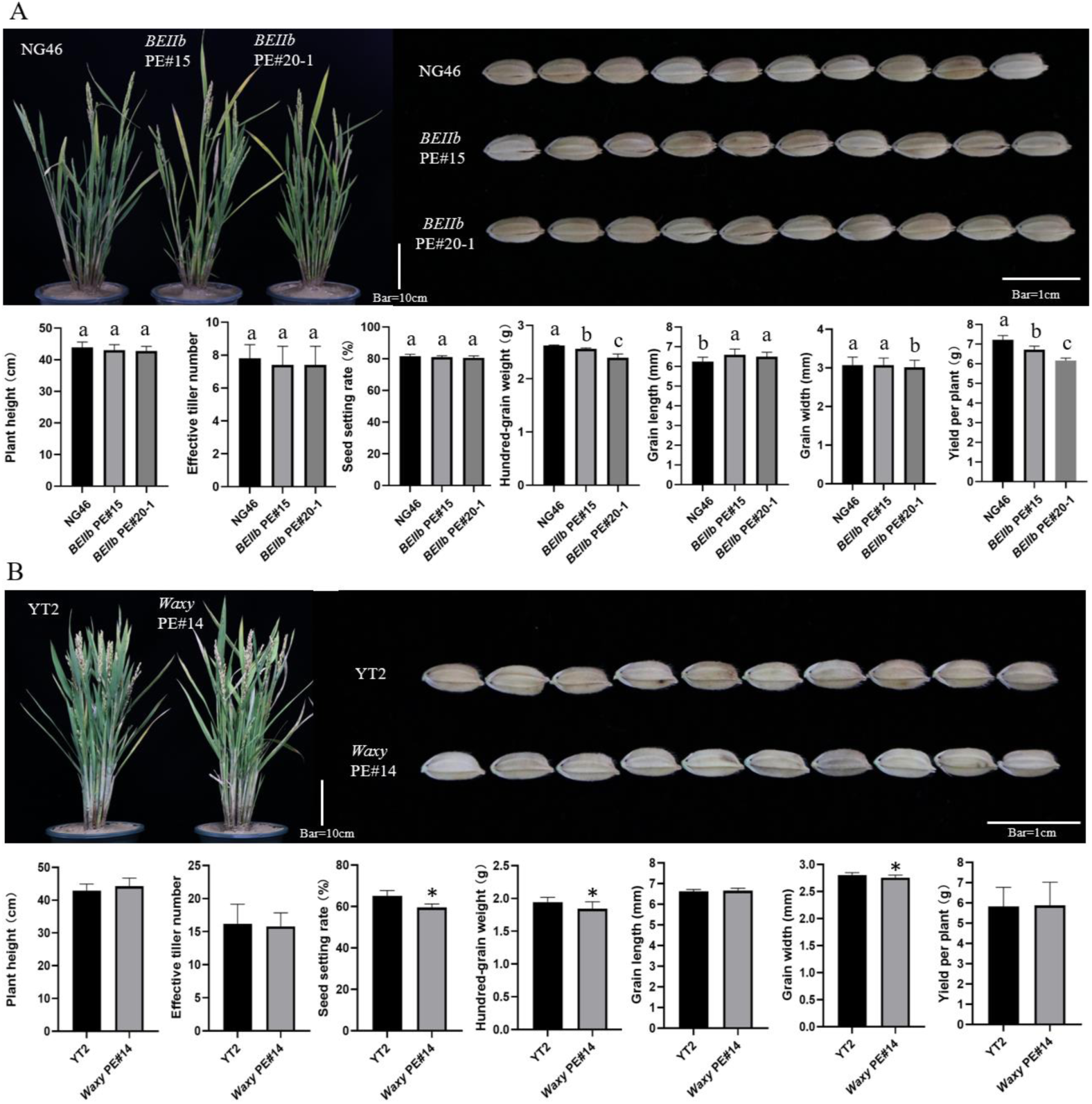
Agronomic traits of selective PE lines including plant architecture at heading stage, grain morphology, plant height, effective tiller number, seed setting rate, hundred-grain weight, grain length, grain width and yield per plant. (A) NG46 and *BEIIb* PE#15, PE#20-1. (B) YT2 and *Wx* PE#14. Bars represent means ± SD (n=5 for main agronomic traits, n=500 for grain shape traits). Student’s t-test; * *P* < 0.05, Different letters above the bars indicate significant differences (*P* < 0.05) determined by one-way analysis of variance followed by LSD.

Interestingly, compared to WT, both *BEIIb* PE#15 and 20-1 showed significantly increased grain length, and PE#20-1 also showed a decrease in grain width. We also measured the *BEIIb* PE#29 for comparison and found that it had significantly increased grain length than NG46 (but significantly shorter than PE#15 and PE#20-1) and decreased grain width (data no showed). Therefore, grain width may be negatively correlated with enhanced RSC. In addition, *Wx* PE#14 also showed a significant decrease in grain width. The above results implied that some pathways for the starch synthesis process and development of grain size might be intertwined in rice.

## 4. Discussion

RS has been found for decades, it is now well-known that foods enriched with RS can prevent obesity, cardiovascular diseases, and help to reduce glycemic and insulin responses, especially for the type 2 diabetic population. Foods with glycemic index (GI) lower than 55 were classified as low-GI foods, and the RSC is a key indicator for measuring GI of any food products. To develop low-GI crop varieties rapidly, the identification and characterization of genes controlling RSC had been actively conducted, it’s now clear that some genes involved in plant starch synthesis play important roles in the generation of RS. The BEIIb was first cloned and identified in the mutant of maize *Amylose*-*Extender* [38]. Subsequently, BEIIb mutants were also found to have high RSC in other crops, including maize, wheat [40], cassava [41], and rice [10–12]. Valid research evidences indicated that loss of function of BEIIb in rice often changes the structure and components of starch in endosperm and then led to being more resistant to pancreatic amylase from animals [42,43]. In wheat, reduced BEIIa expression, a major active BEII enzyme in endosperm, resulted in high-amylose grain, and feeding experiments showed improved indices of gastrointestinal health in Rats [44]. Another well-known example to the public is the naturally occurring wrinkled peas, a mutant of starch branching enzyme I gene, which showed altered starch assembly and improved glucose homeostasis in humans [45].

However, many of the single or double gene mutants enriched with RS in endosperm, also have much higher AC(≥20%). This problem is prominent especially in rice, since high AC always corresponds to lower taste quality and consumer acceptance. Therefore, in this study, we used PE and attempted two strategies to decrease the AC for the high RS lines produced by manipulating *BEIIb*. For *BEIIb* PE, 6gRNAs were designed to target the core region of promoter to produce novel alleles that may downregulate *BEIIb* expression to an appropriate extent. After screening and sequencing of the homozygous offspring, we found a high rate of large fragment deletion in *BEIIb* PE lines. In the same way, 6gRNAs targeted the promoter of *Wx* in a variety with *be2b* loss of function mutation (YT2) to decrease the AC. Interestingly, the rate of large fragment deletion was low, probably due to the relatively long distance between the target sites. Previous studies showed that large deletions or rearrangements usually cause effective phenotypic changes [22, 46]. Similar results were found in this study. All the selected PE-lines including five lines from *BEIIb* and one line from *Wx* displayed large deletions between the target sites, the most extreme example is the *BEIIb* PE#22, which showed almost loss of 2kb promoter, despite that, the RS content of this line was not the highest, suggesting the potentially recruiting of other regulatory elements from upstream sequence. Large deletions would be more effective due to the high chance to completely remove *cis*-regulatory elements which cannot always be precisely predicted bioinformatically. Finally, we obtained 2 BEIIb PE lines and one *Wx* PE line with high RS content (≥5%) but relatively lower AC (≤20%). The taste quality of the two lines was acceptable and within the range of high-easting quality varieties, although it significantly deteriorated compared to WT (NG46). The results highlighted that importance of choosing the background variety for gene editing and the technique adopted. Although *Wx* PE#14 line had much lower AC and better taste quality compared to the popular low-GI (high-RS) variety YT2. However, its RS content was significantly elevated as well. From this perspective downregulating the expression of *Wx* gene seems to be another good strategy to significantly improve the taste quality of high-RS rice varieties based on *be2b* already on the market.

Our analysis of 3 *BEIIb* PE lines showed that deletions of T2 to T6 region of *BEIIb* promoter significantly influence its expression level in endosperm. This region was predicted to have binding sites for many TFs, we then subsequently discovered some of TFs were among the coexpression gene set of *BEIIb* and/or within the important TFs reported for *BEIIIb*. Dual-luciferase reporter assay further confirmed that *OsbZIP60*, *OsMYB3*, *OsRSL2*, and *OsERF44* can activate the expression of *BEIIIb* target promoter region in rice protoplasts, though the fold change is not high, further efforts may be invested into searching for more key TFs or try multiple-gene knock out for these TFs to confirm their relationships with RS production in rice.

Increasing RS content is usually accompanied by increased AC based on previous studies. In this study, we also observed a strong and positive correlation between RSC and AC using the *BEIIb* PE lines. Intriguingly, in the *Wx* PE lines, the correlation between RSC and AC seems to be negative. This data created a confusing scene to understand about the real relationship between AC and RSC in the defective *BEIIb* background.

Previous studies showed that loss function of *BEIIb* mutant (*be2b*) has amylopectin structure with enriched long chains, and its starch crystalline exhibited a B-type pattern in contrast to the typical A-type pattern found in the WT [47, 48]. We also detected the crystalline pattern of some *BEIIb* PE lines through the XRD diffraction technique, and found that the starch crystalline of all lines except *BEIIb* PE#22 with the lowest RSC was B-type pattern (Table S3 and SP Fig.2). B-type starch granules are thought to be more resistant to enzyme hydrolysis due to the compact structure, which limits the accessibility of digestive enzymes and displays a nonporous, smooth surface [48, 49]. A typical example is the natural starch from raw potatoes that is mainly enriched with RS2 has the B-type crystallinity but is heat-sensitive with drastically reduced RSC during the conventional cooking process [49, 50]. Further analysis of the starch structure from the rice *wx ae* double-mutant showed an almost identical B-type crystallinity pattern to the *ae* mutant starch [43]. More importantly, despite the absence of amylose, *wx ae* mutant also showed high RSC in raw starch [51, 52]. Thus, amylose in endosperm is not required for the production of RS2 starch with B-type crystallinity. Based on previous findings, different high RS mutants produce different types of RS. For example, the double mutant of *SSIIIa* and *SSIIIb* produced abundant heat-generated RS during cooking, whereas its raw RS was about one-third of its cooked product [18]. In contrast, the RS of *be2b* mutant rice flour was also cooking-sensitive resemble the starch from raw potatoes and had about three times higher RSC in raw flour than its cooked flour [16, 42]. Therefore, we speculated that the *be2b* mutant and our PE lines with downregulated expression of BEIIb predominantly produce the RS2 with B-type crystallinity, which did not participation of amylose.

The downregulation of *BEIIb* expression in its PE lines decreased the production of short chains in amylopectin, but it also promoted the expression of *Wx* and other starch synthases used the redundant glycose substrates to produce more amylose or elongating short chains in amylopectin, the amylopectin enrich with long chains further promote the transformation of starch crystallinity from A type to B type, which finally led to production of RS2 (Fig. 7). Notably, the change of this structure for amylopectin significantly reduces the number of exposed non-reducing ends, which is also disadvantageous for amylase to hydrolyze starch [53]. On the contrary, when decreased expression of *Wx* in its PE lines in the *be2b* background, the redundant glycose substrates could not be fully utilized by the downregulated *Wx* gene and more substrates are available for other enzymes responsible for the elongation of short branches in amylopectin. As a consequence, more RS2 was produced in *Wx* PE lines. We also provided a convincing data from chain length distribution analysis of amylopectin for these PE lines, especially for *BEIIb* PE lines, which their RSC was significantly and positively correlated with the total amount of long branch chains in amylopectin. (Fig. 5C). Indeed, the correlation was slightly stronger than that between RSC and AC even taken the data of WT into calculation. Similar result had been reported by analyzing the percentage of amylopectin long chains between *be2b*, *ss3a*/*be2b*, and *ss1*/*be2b* mutants [37]. Taken together, these results provided reasonable explanations for the confusing relationships between RSC and AC in our different PE lines.

**Fig.7.**
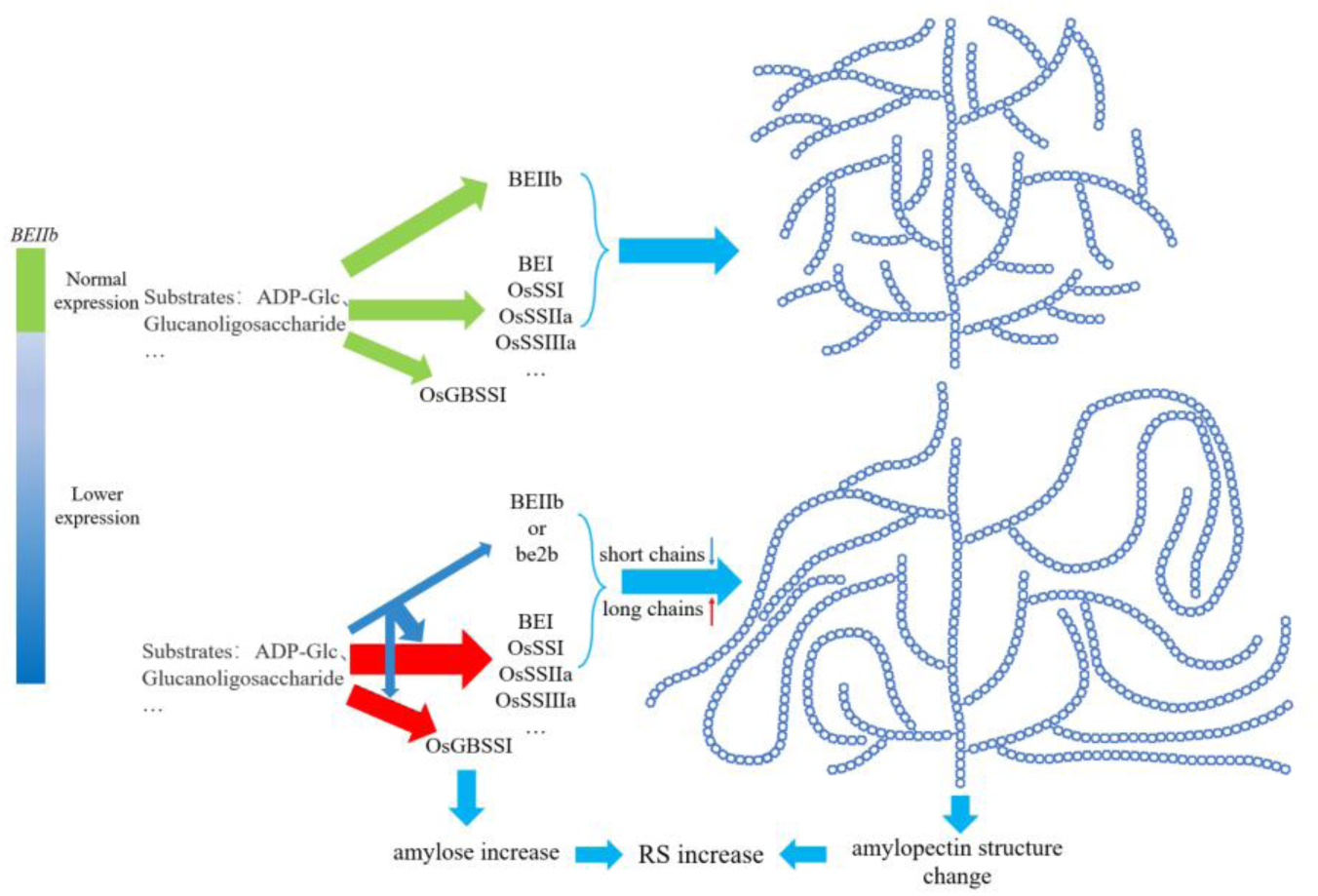
Schematic diagram of RS production in rice *BEIIb* deficiency material.

Nevertheless, even though it is very likely that the starch of *be2b* was enriched with RS2, we could not completely exclude that other types of RS (heat-insensitive or heat-produced) could be produced in *be2b* mutant. Besides, the *be2b* mutant starch also had a stronger retrogradation ability [43]. Moreover, compared to the gelatinized rice flour, the mashed cooked rice that is close to human eating custom still maintained about 90% RSC of raw rice flour in *be2b* background based on previous study [16]. Therefore, further characterization of RS produced by our PE lines in this study is required to facilitate its use in manufacturing different property of RS enriched foods.

## 5. Conclusion

We obtained two *BEIIb* PE lines and one *Wx* PE line with high RSC (≥5%) but relatively lower AC (≤20%), demonstrating that PE of BEIIb or *Wx* genes have the potential to generate germplasms with acceptable AC and RSC. These PE lines did not show significant changes in most agronomic traits, only the BEIIb PE lines had slightly reduced yield compared to the WT, so they have the potential to be used in agricultural production. Although AC increases with increase of RSC in the BEIIb PE population, its contribution to RS production should be further investigated, since the amount of long-chain branches of amylopectin was also high positively correlated with RSC and increased production of RS was previously reported for *wx ae* double-mutant (without AC). In addition, the fact that our *Wx* PE lines in *be2b* mutant background had reduced AC but increased RSC also suggested that the relationship between AC and RSC is complex and need to be further studied.

## Data availability

All materials created in this study will be made available on request to corresponding author.

## CRediT authorship contribution statement

**Yuelong Lin:** Writing–original draft, Investigation, Formal analysis, Data curation. **Qianqi Guo:** Methodology, Investigation, Validation. **XingLi Xu:** Investigation, Validation. **Huaiying Gu:** Investigation, Validation. **MingHong Hu:** Investigation, Validation. **Weiying Kong:** Investigation, Validation. **Yongjun Wu**: Supervision, Resources, Funding acquisition, **Lijun Meng**: Writing–review & editing, Resources, Funding acquisition, Supervision, **Guoyou Ye**: Writing–review & editing, Supervision, Resources, Funding acquisition, Conceptualization.

## Acknowledgments

This study was supported by The Special Financial Fund of Foshan in 2023–Cooperation project for high-level agricultural science and technology demonstration city construction in Guangdong Province.

## Appendix A. Supplementary data

## Supplementary data

**Supplementary Fig.1.**
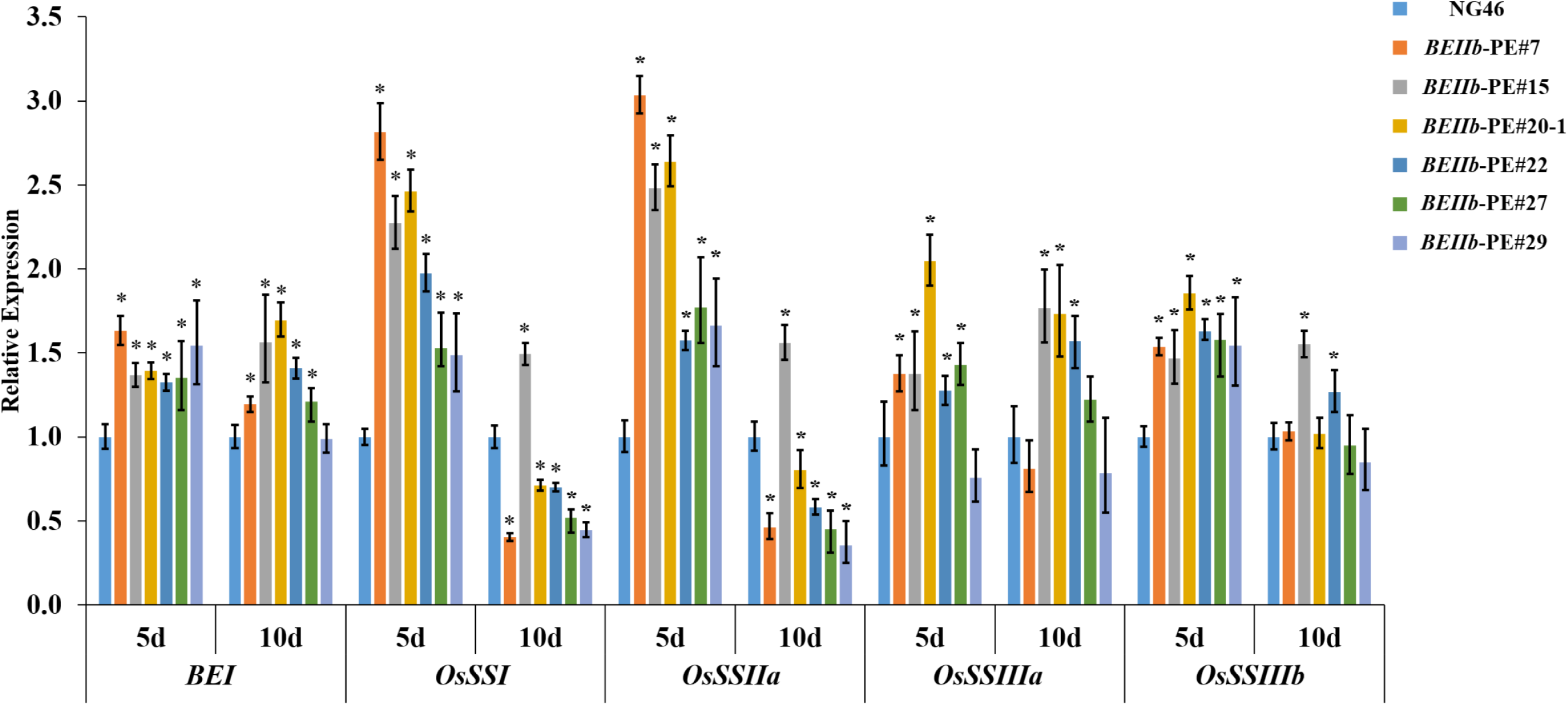
Expression analysis of major enzymes in the starch biosynthesis pathway of endosperm for *BEIIb* PE lines. Bars represent means ± SD, Student’s t-test; * *P* < 0.05

**Supplementary Fig.2.**
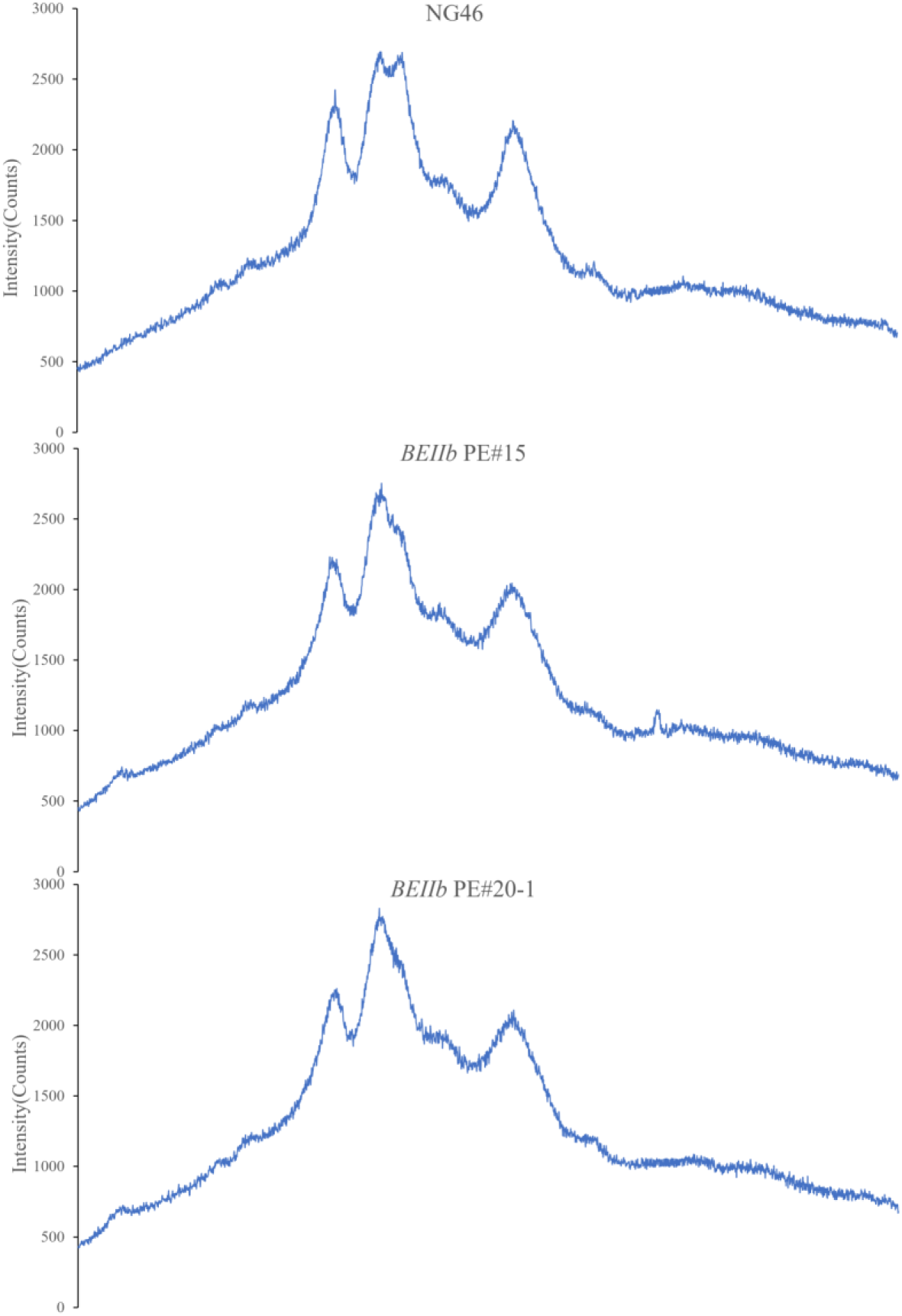
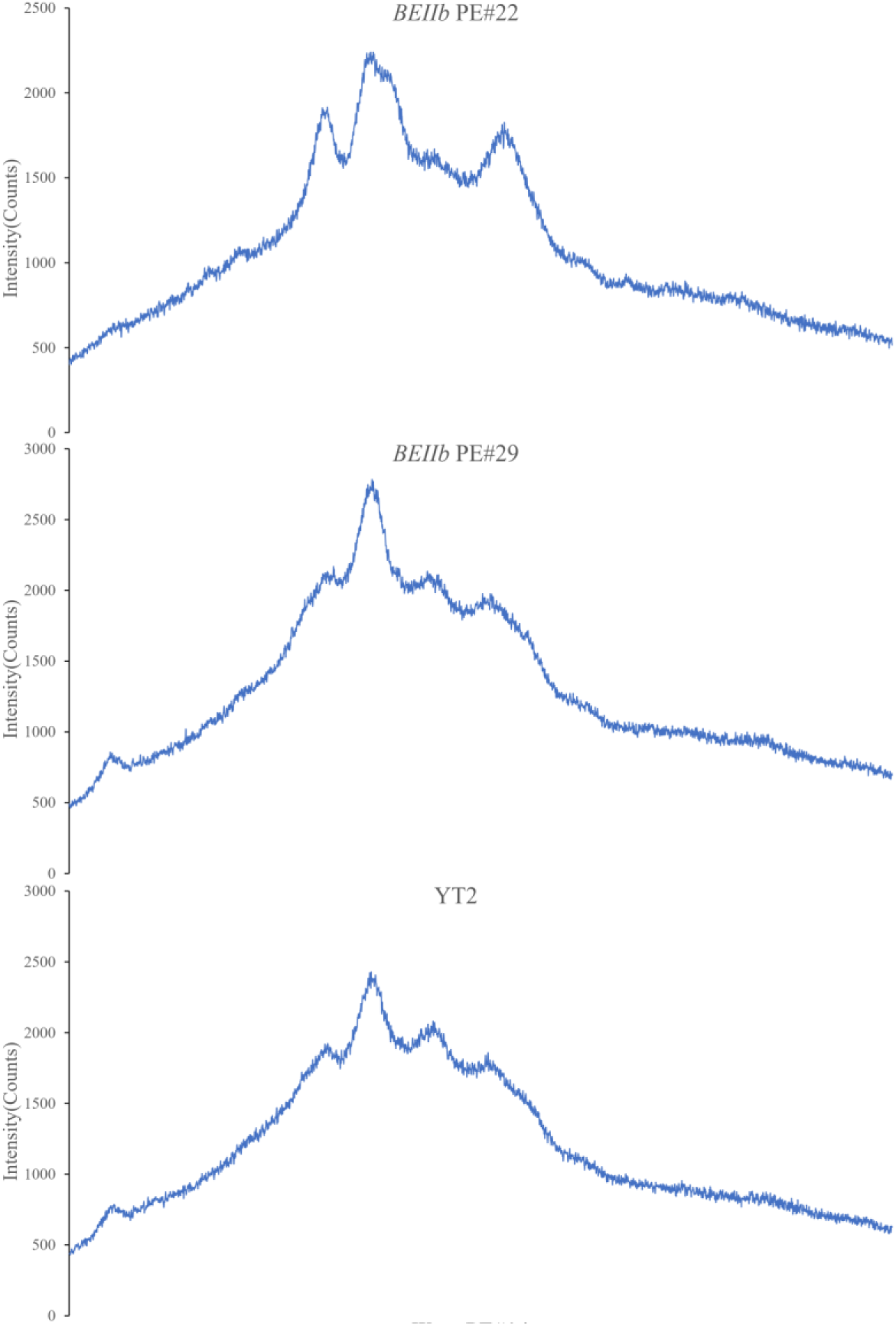
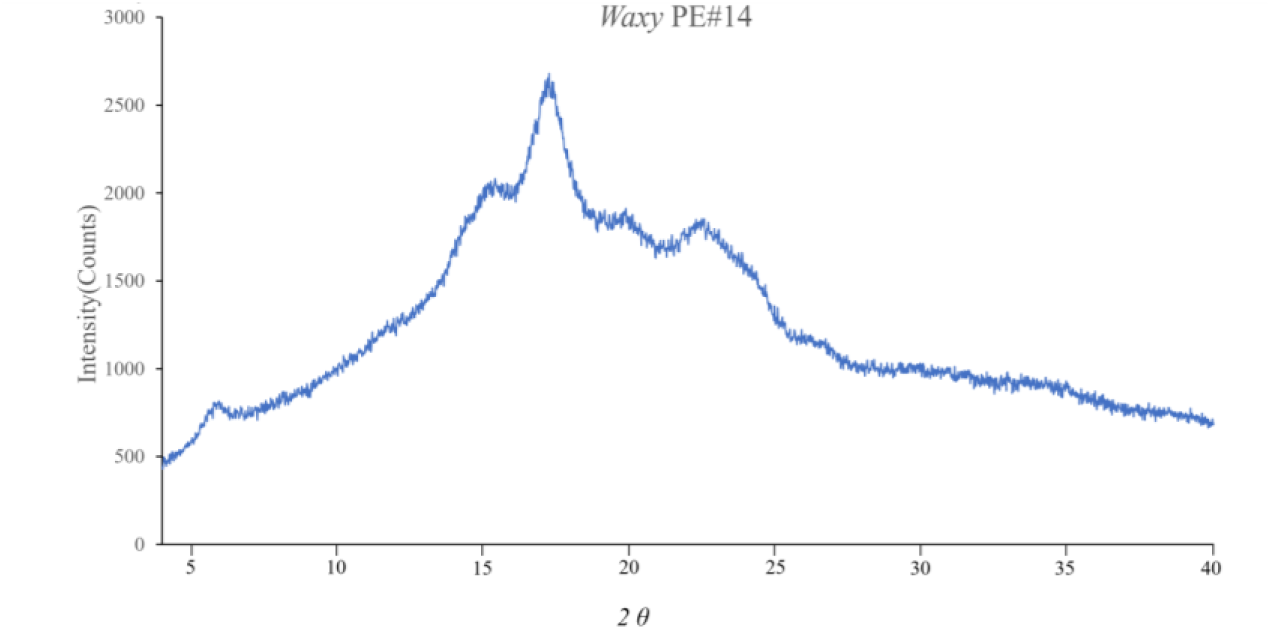
X-ray diffraction analysis of starch granules for some PE lines

**Table S1.**
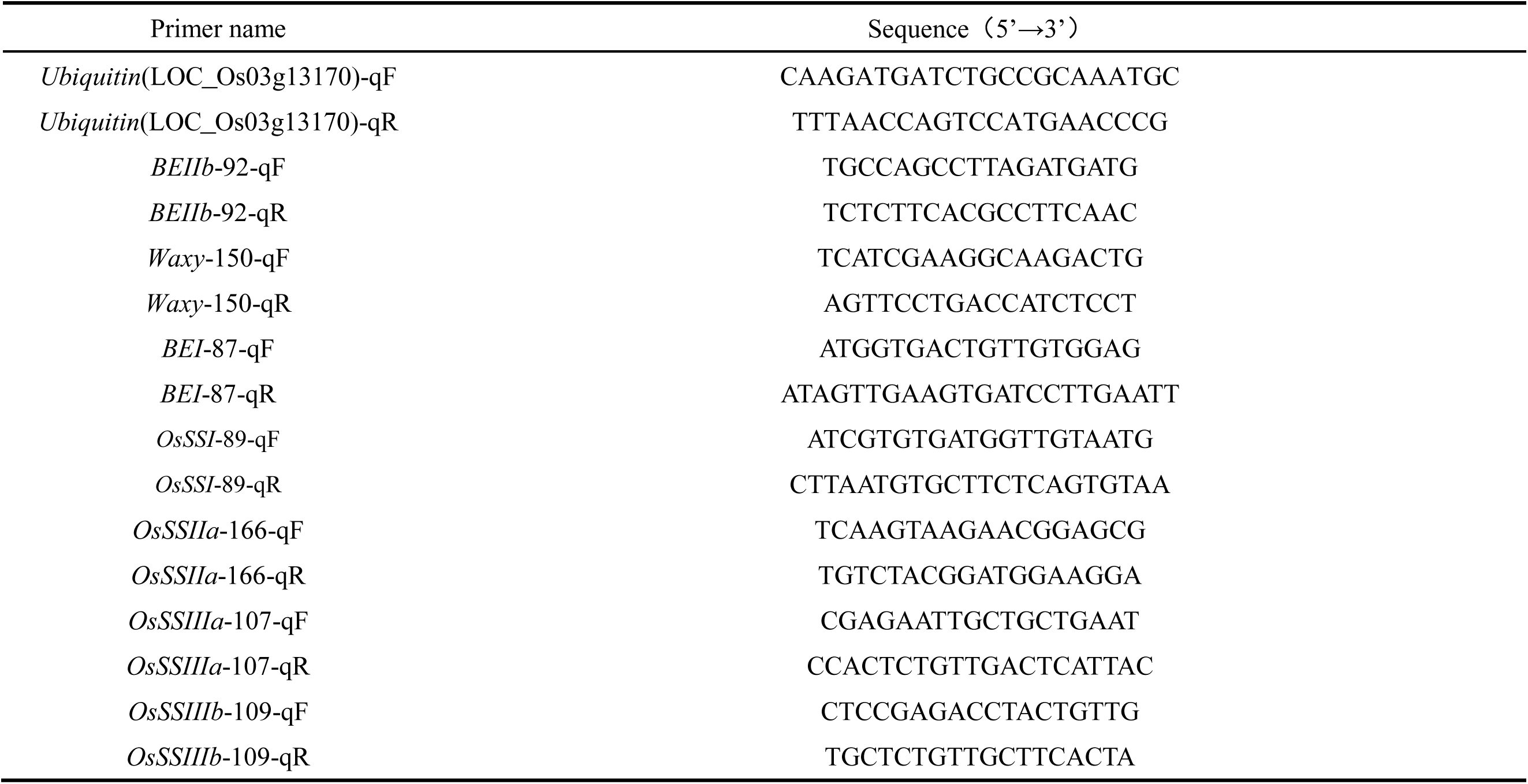
The primers used for RT-qPCR in this study.

**Table S2.**
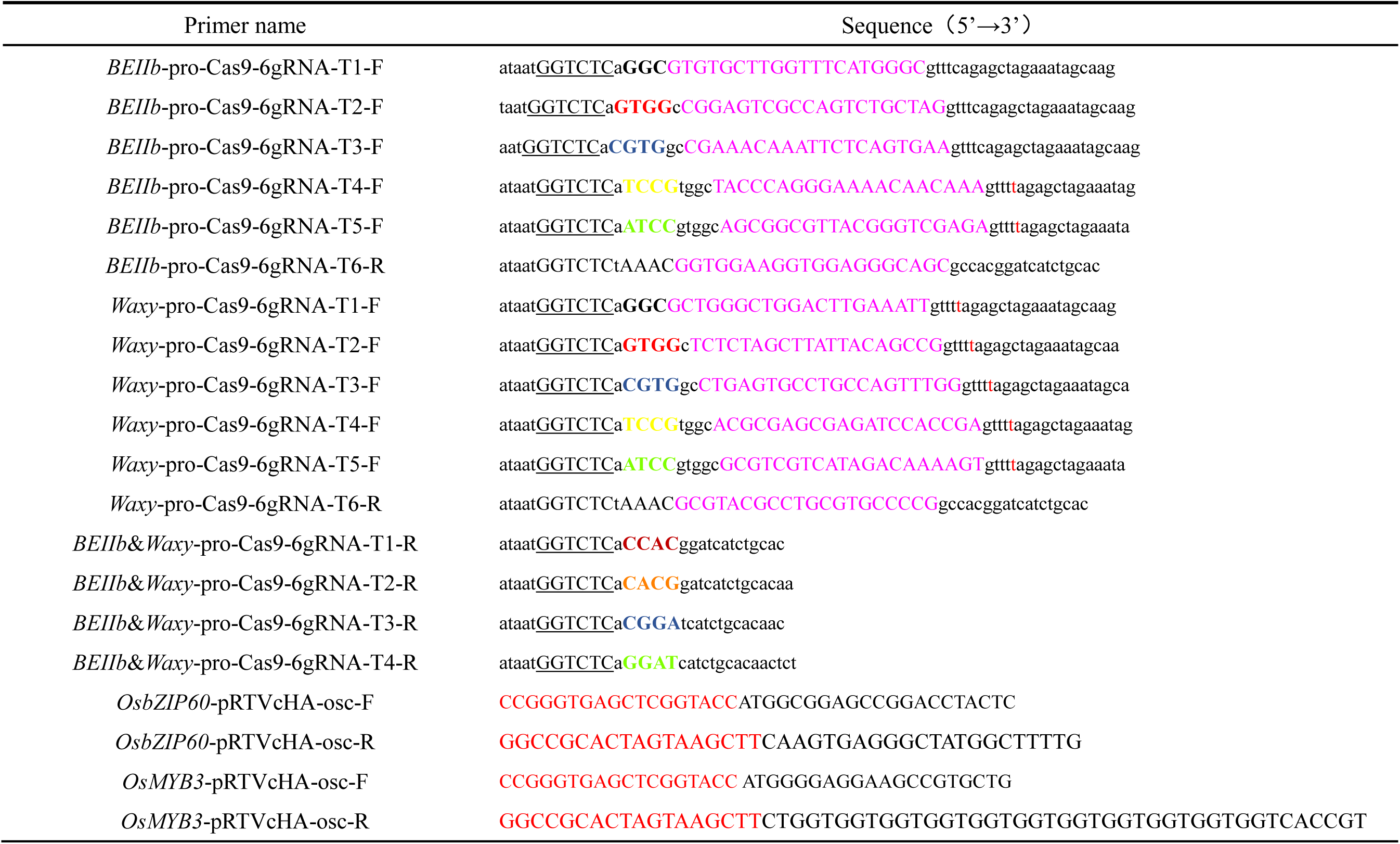
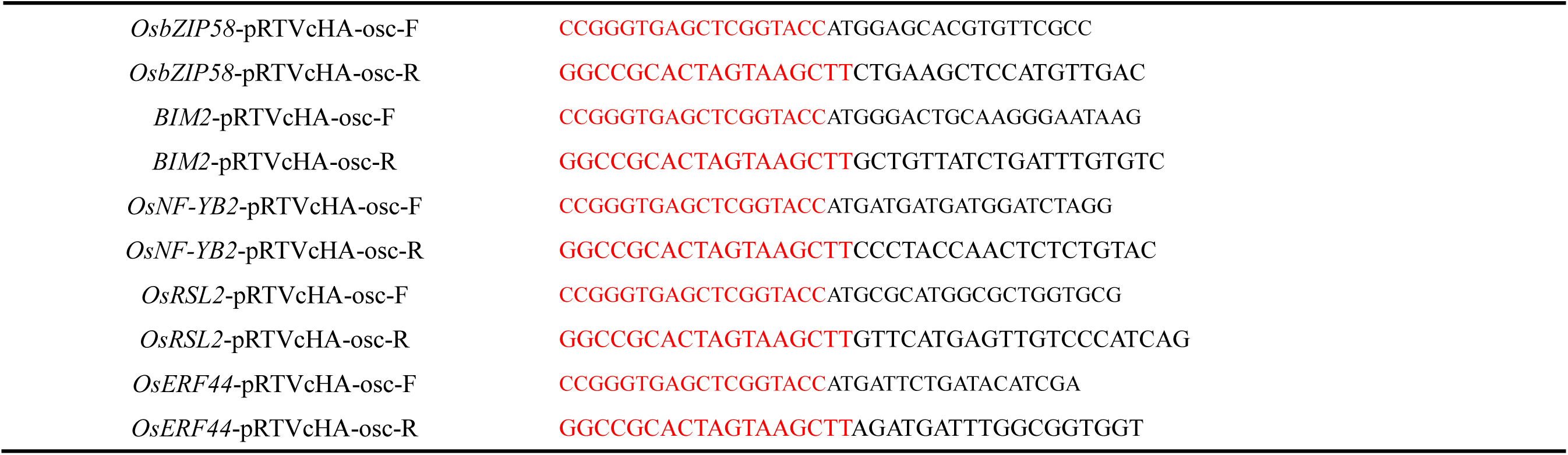
The primers used for vector construction in this study.

**Table S3.**
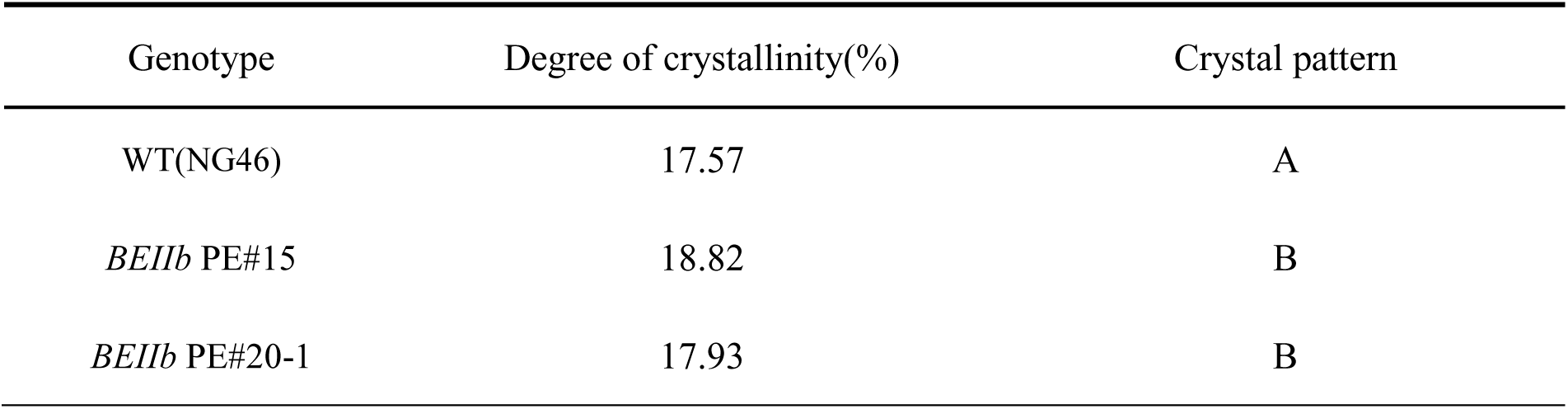
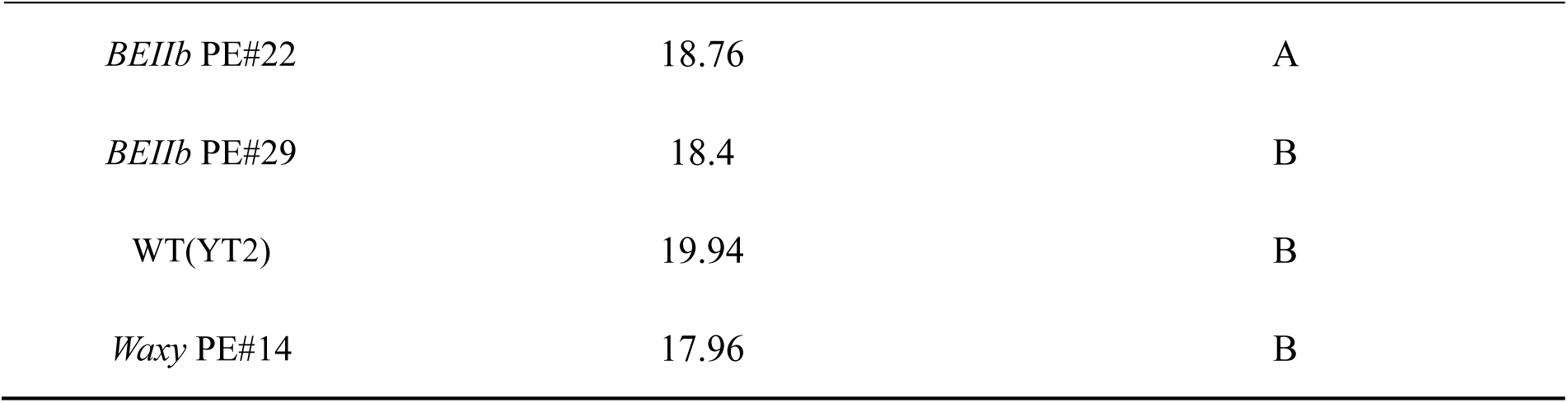
Analysis results of XRD for some PE lines.

