## Supplemental data for "Promoter editing of starch branching enzyme IIb and granule-bound starch synthase I balances resistant starch content and amylose content in rice"

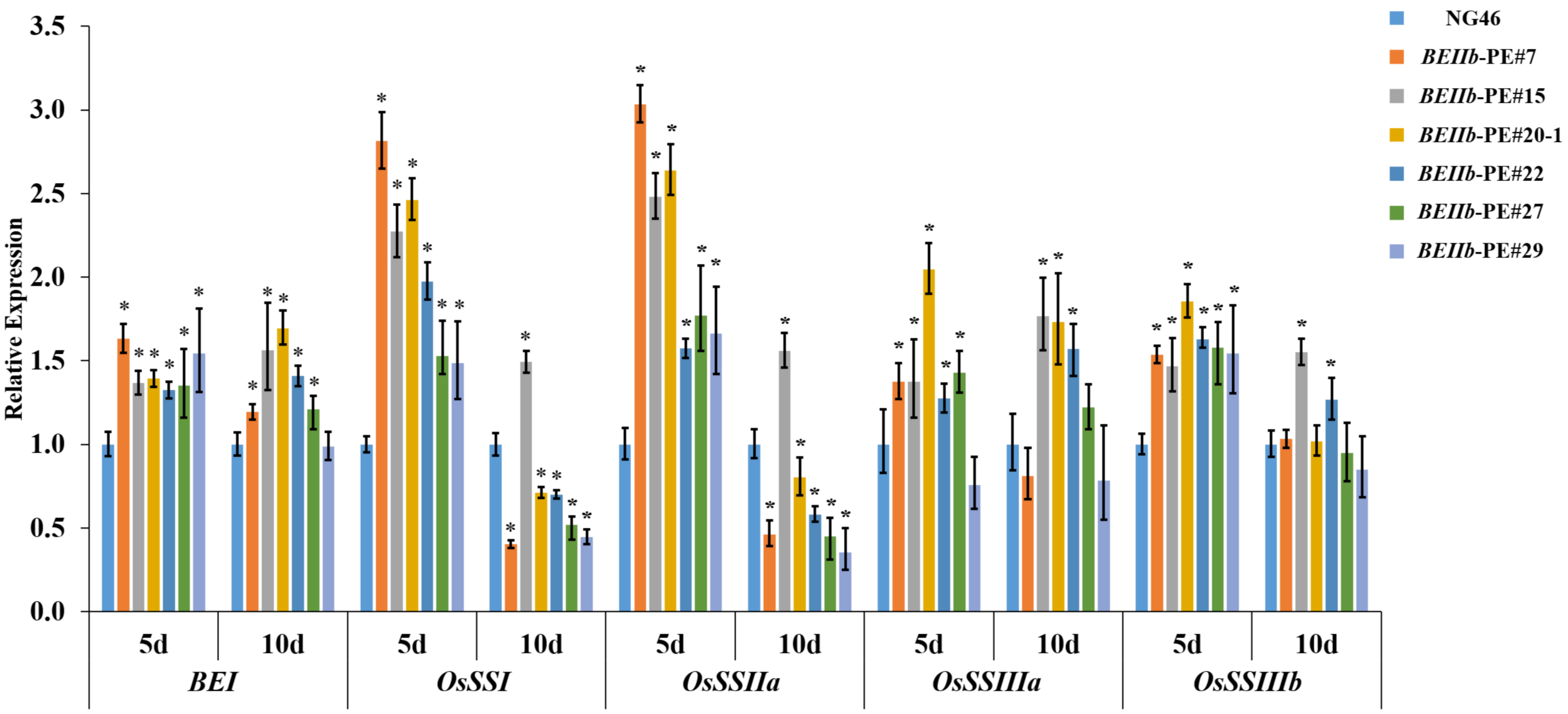


Supplementary Fig.1 Expression analysis of major enzymes in the starch biosynthesis pathway of endosperm for *BEIIb* PE lines. Bars represent means ± SD, Student's t-test; * $P < 0.05$

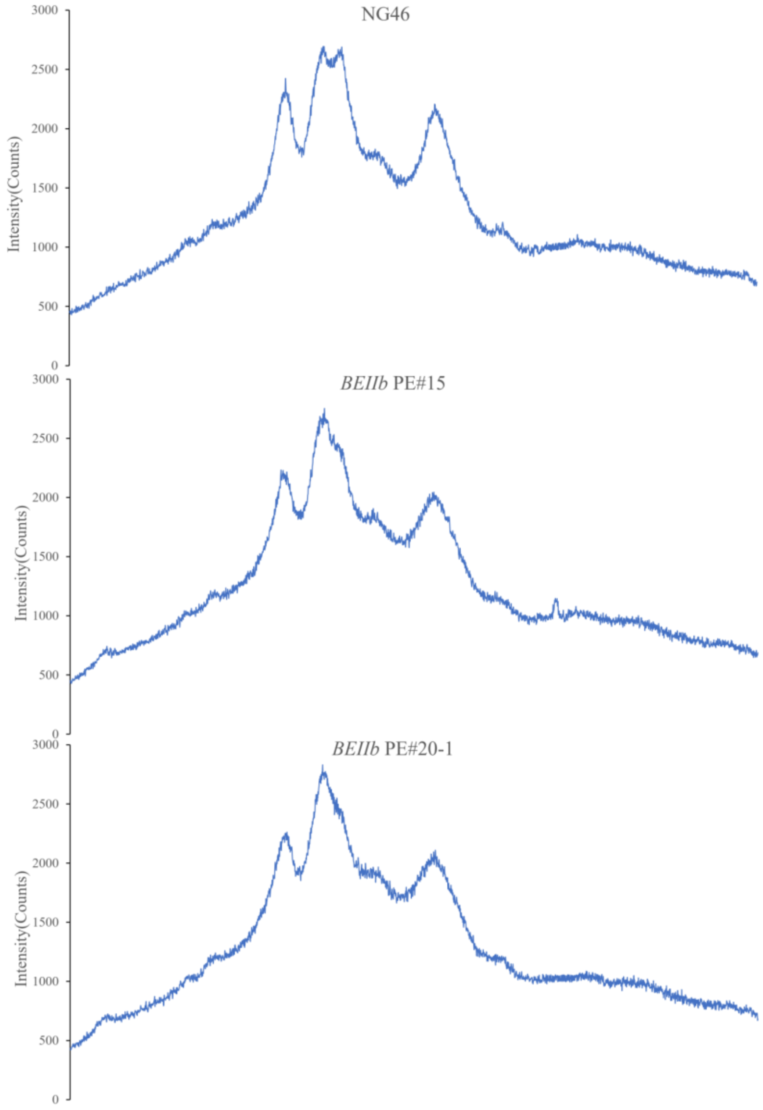
NG46
*BEIIb* PE#15
*BEIIb* PE#20-1
Intensity(Counts)
3000
2500
2000
1500
1000
500
0

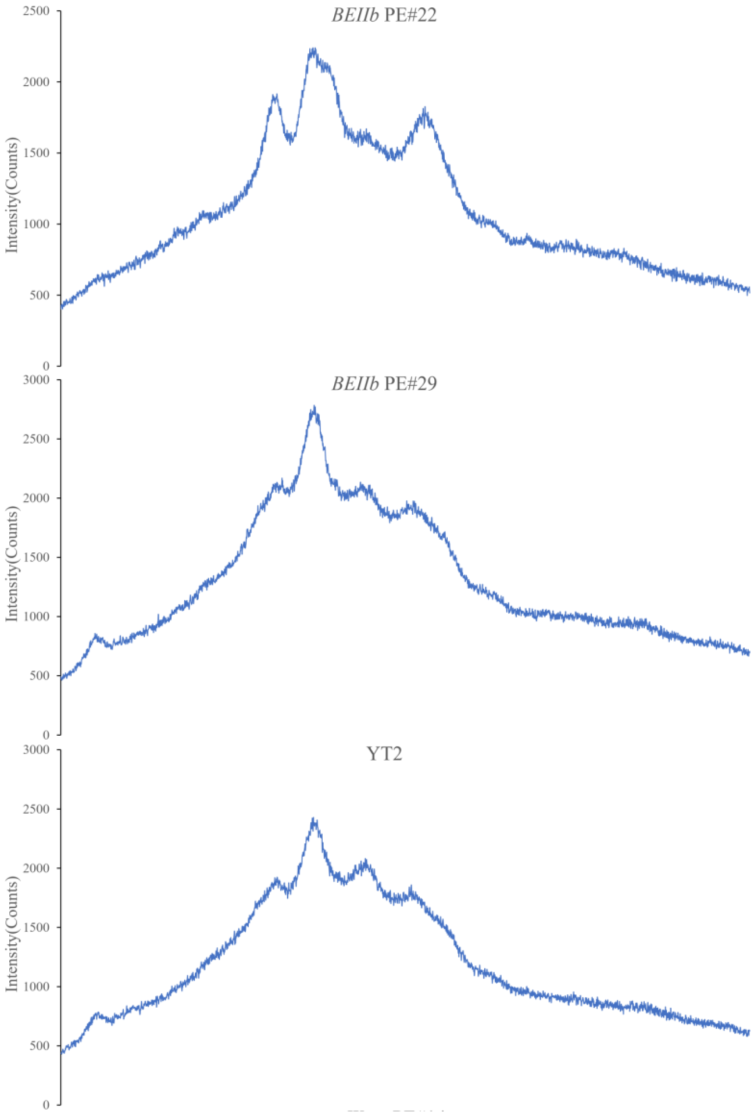
*BEIIb* PE#22
Intensity(Counts)
2500
2000
1500
1000
500
0
*BEIIb* PE#29
Intensity(Counts)
3000
2500
2000
1500
1000
500
0
YT2
Intensity(Counts)
3000
2500
2000
1500
1000
500
0

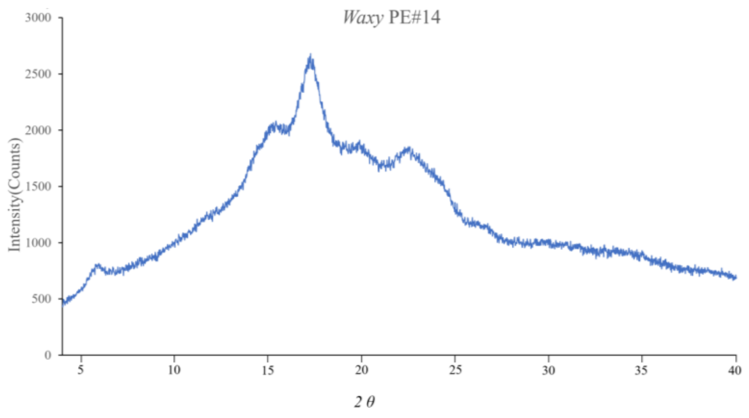


Supplementary Fig.2 X-ray diffraction analysis of starch granules for some PE lines

Table S1. The primers used for RT-qPCR in this study

| Primer name | Sequence（5’→3’） |
|---|---|
| *Ubiquitin*(LOC_Os03g13170)-qF | CAAGATGATCTGCCGCAAATGC |
| *Ubiquitin*(LOC_Os03g13170)-qR | TTTAACCAGTCCATGAACCCG |
| *BEIIb*-92-qF | TGCCAGCCTTAGATGATG |
| *BEIIb*-92-qR | TCTCTTCACGCCTTCAAC |
| *Waxy*-150-qF | TCATCGAAGGCAAGACTG |
| *Waxy*-150-qR | AGTTCCTGACCATCTCCT |
| *BEI*-87-qF | ATGGTGACTGTTGTGGAG |
| *BEI*-87-qR | ATAGTTGAAGTGATCCTTGAATT |
| *OsSSI*-89-qF | ATCGTGTGATGGTTGTAATG |
| *OsSSI*-89-qR | CTTAATGTGCTTCTCAGTGTAA |
| *OsSSIIa*-166-qF | TCAAGTAAGAACGGAGCG |
| *OsSSIIa*-166-qR | TGTCTACGGATGGAAGGA |
| *OsSSIIIa*-107-qF | CGAGAATTGCTGCTGAAT |
| *OsSSIIIa*-107-qR | CCACTCTGTTGACTCATTAC |
| *OsSSIIIb*-109-qF | CTCCGAGACCTACTGTTG |
| *OsSSIIIb*-109-qR | TGCTCTGTTGCTTCACTA |

Table S2. The primers used for vector construction in this study

| Primer name | Sequence（5’→3’） |
|---|---|
| *BEIIb*-pro-Cas9-6gRNA-T1-F | ataatGGTCTCa**GGC**GTGTGCTTGGTTTCATGGGCgtttcagagctagaaatagcaag |
| *BEIIb*-pro-Cas9-6gRNA-T2-F | taatGGTCTCa**GTGG**cCGGAGTCGCCAGTCTGCTAGgtttcagagctagaaatagcaag |
| *BEIIb*-pro-Cas9-6gRNA-T3-F | aatGGTCTCa**CGTG**gcCGAAACAAATTCTCAGTGAAgtttcagagctagaaatagcaag |
| *BEIIb*-pro-Cas9-6gRNA-T4-F | ataatGGTCTCaTCCGtggcTACCCAGGGAAAACAACAAAgttttagagctagaaatag |
| *BEIIb*-pro-Cas9-6gRNA-T5-F | ataatGGTCTCaATCCgtggcAGCGGCGTTACGGGTCGAGAgttttagagctagaaata |
| *BEIIb*-pro-Cas9-6gRNA-T6-R | ataatGGTCTCtAAACGGTGGAAGGTGGAGGGCAGCgccacggatcatctgcac |
| *Waxy*-pro-Cas9-6gRNA-T1-F | ataatGGTCTCa**GGC**GCTGGGCTGGACTTGAAATTgttttagagctagaaatagcaag |
| *Waxy*-pro-Cas9-6gRNA-T2-F | ataatGGTCTCa**GTGG**cTCTCTAGCTTATTACAGCCGgttttagagctagaaatagcaa |
| *Waxy*-pro-Cas9-6gRNA-T3-F | ataatGGTCTCa**CGTG**gcCTGAGTGCCTGCCAGTTTGGgttttagagctagaaatagca |
| *Waxy*-pro-Cas9-6gRNA-T4-F | ataatGGTCTCaTCCGtggcACGCGAGCGAGATCCACCGAgttttagagctagaaatag |
| *Waxy*-pro-Cas9-6gRNA-T5-F | ataatGGTCTCaATCCgtggcGCGTCGTCATAGACAAAAGTgttttagagctagaaata |
| *Waxy*-pro-Cas9-6gRNA-T6-R | ataatGGTCTCtAAACGCGTACGCCTGCGTGCCCCGgccacggatcatctgcac |
| *BEIIb*&*Waxy*-pro-Cas9-6gRNA-T1-R | ataatGGTCTCa**CCAC**ggatcatctgcac |
| *BEIIb*&*Waxy*-pro-Cas9-6gRNA-T2-R | ataatGGTCTCaCACGgatcatctgcacaa |
| *BEIIb*&*Waxy*-pro-Cas9-6gRNA-T3-R | ataatGGTCTCa**CGGA**tcatctgcacaac |
| *BEIIb*&*Waxy*-pro-Cas9-6gRNA-T4-R | ataatGGTCTCaGGATcatctgcacaactct |
| *OsbZIP60*-pRTVcHA-osc-F | CCGGGTGAGCTCGGTACCATGGCGGAGCCGGACCTACTC |
| *OsbZIP60*-pRTVcHA-osc-R | GGCCGCACTAGTAAGCTTCAAGTGAGGGCTATGGCTTTTG |
| *OsMYB3*-pRTVcHA-osc-F | CCGGGTGAGCTCGGTACC ATGGGGAGGAAGCCGTGCTG |
| *OsMYB3*-pRTVcHA-osc-R | GGCCGCACTAGTAAGCTTCTGGTGGTGGTGGTGGTGGTGGTGGTGGTGGTCACCGT |

| | |
|---|---|
| *OsbZIP58*-pRTVcHA-osc-F | CCGGGTGAGCTCGGTACCATGGAGCACGTGTTCGCC |
| *OsbZIP58*-pRTVcHA-osc-R | GGCCGCACTAGTAAGCTTCTGAAGCTCCATGTTGAC |
| *BIM2*-pRTVcHA-osc-F | CCGGGTGAGCTCGGTACCATGGGACTGCAAGGGAATAAG |
| *BIM2*-pRTVcHA-osc-R | GGCCGCACTAGTAAGCTTGCTGTTATCTGATTTGTGTC |
| *OsNF-YB2*-pRTVcHA-osc-F | CCGGGTGAGCTCGGTACCATGATGATGATGGATCTAGG |
| *OsNF-YB2*-pRTVcHA-osc-R | GGCCGCACTAGTAAGCTTCCCTACCAACTCTCTGTAC |
| *OsRSL2*-pRTVcHA-osc-F | CCGGGTGAGCTCGGTACCATGCGCATGGCGCTGGTGCG |
| *OsRSL2*-pRTVcHA-osc-R | GGCCGCACTAGTAAGCTTGTTCATGAGTTGTCCCATCAG |
| *OsERF44*-pRTVcHA-osc-F | CCGGGTGAGCTCGGTACCATGATTCTGATACATCGA |
| *OsERF44*-pRTVcHA-osc-R | GGCCGCACTAGTAAGCTTAGATGATTTGGCGGTGGT |

Table S3. Analysis results of XRD for some PE lines

| Genotype | Degree of crystallinity(%) | Crystal pattern |
|---|---|---|
| WT(NG46) | 17.57 | A |
| *BEIIb* PE#15 | 18.82 | B |
| *BEIIb* PE#20-1 | 17.93 | B |

| | | |
|---|---|---|
| *BEIIb* PE#22 | 18.76 | A |
| *BEIIb* PE#29 | 18.4 | B |
| WT(YT2) | 19.94 | B |
| *Waxy* PE#14 | 17.96 | B |
